# Pairwise Relative Distance (PRED) is an intuitive and robust metric for assessing vector similarity and class separability

**DOI:** 10.1101/2021.08.13.456194

**Authors:** Aarush Mohit Mittal, Andrew C. Lin, Nitin Gupta

## Abstract

Scientific studies often require assessment of similarity between ordered sets of values. Each set, containing one value for every dimension or class of data, can be conveniently represented as a vector. The commonly used metrics for vector similarity include angle-based metrics, such as cosine similarity or Pearson correlation, which compare the relative patterns of values, and distance-based metrics, such as the Euclidean distance, which compare the magnitudes of values. Here we evaluate a newly proposed metric, pairwise relative distance (PRED), which considers both relative patterns and magnitudes to provide a single measure of vector similarity. PRED essentially reveals whether the vectors are so similar that their values across the classes are separable. By comparing PRED to other common metrics in a variety of applications, we show that PRED provides a stable chance level irrespective of the number of classes, is invariant to global translation and scaling operations on data, has high dynamic range and low variability in handling noisy data, and can handle multi-dimensional data, as in the case of vectors containing temporal or population responses for each class. We also found that PRED can be adapted to function as a reliable metric of class separability even for datasets that lack the vector structure and simply contain multiple values for each class.

## Introduction

Vectors are ubiquitous data structures. As a result, the assessment of vector similarity is one of the most frequently performed data operations in diverse areas of science and engineering. To list examples within only biology, vector similarity has been used to show that reef fish species in different ecoregions resemble each other in traits, not taxonomy or phylogeny (McLean et al., 2021); that cancerous cell lines’ gene expression patterns cluster according to their tissue of origin and cancer stage (Ross et al., 2000); and that certain brain regions have similar fMRI brain activation patterns over time, suggesting they are functionally connected (Sasai et al., 2021). In these examples, the vectors represented the trait, taxonomical or phylogenetic properties of each ecoregion; the gene expression profile of each cell line; and the temporal activation pattern of each brain region, respectively. Similarly, other examples of scientific data that can be represented as vectors include the firing rates of a cortical neuron to different visual stimuli (Hubel and Wiesel, 1962; Stringer et al., 2019; Victor and Purpura, 1996), the eye blinking rates of a human under different airflow conditions (VanderWerf et al., 2003), and the sensory preferences of an animal to a given stimulus at different time points (Buchanan et al., 2015; Honegger et al., 2020; Kain et al., 2015; Linneweber et al., 2020)). Any scientific question involving the comparison of such vectors requires metrics that can determine the level of similarity between vectors.

Common metrics for vector similarity include Pearson’s correlation, cosine similarity, and Euclidean distance. Distance-based metrics, like Euclidean distance or Manhattan distance, compare the magnitude of difference between the values in the two vectors. On the other hand, angle-based metrics, like the cosine similarity or the Pearson’s correlation, compare the relative pattern of values within a vector with that in another vector. To take a straightforward example, consider the vectors [1 2 3] and [10 20 30]. A distance-based metric would call them different, while an angle-based metric would call them very similar. On the other hand, the vectors [1 2 3] and [3 2 1] would be described as relatively similar by the distance-based metrics and dissimilar by the angle-based metrics. Both types of metrics provide useful and complementary information; however, in practice, multiple metrics are rarely used together. In many applications, instead of choosing between one of the two types of metrics, it would be desirable to combine the similarity in the magnitudes and the similarity in the relative patterns into a single, reliable indicator of vector similarity.

We recently devised a metric, called Pairwise Relative Distance (PRED), to quantify the level of similarity in different individuals’ neuronal responses to the same set of odors (Mittal et al., 2020). PRED captured the similarities both in the absolute values and the across-odor patterns of the responses and provided more intuitive values of similarity than correlation in quantifying stereotypy in sensory responses (Mittal et al., 2020). These initial results led us to ask whether PRED could serve as a general-purpose metric for analyzing vector similarity in different types of datasets.

Here, we generalize PRED as a robust metric for assessing vector similarity and class separability. Using simulations and experimental data, we show the advantages of PRED over the commonly used metrics and demonstrate its reliability in analyzing noisy or incomplete data. We illustrate PRED’s ability to capture the similarity in temporal or population-level data while preserving the dataset’s structure. Although we illustrate the usefulness of PRED using examples from the olfactory system, one can use PRED equally well in other sensory modalities in neuroscience, non-neuroscience biological fields like the examples described above, and non-biological fields like machine learning. Overall, our results present Pairwise Relative Distance as a reliable metric of similarity or separability in neuroscience and beyond.

## Results

### PRED as a general metric for vector similarity

In this work, we generalize PRED to all datasets that can be expressed as a matrix, whose columns are specific classes (dimensions) and rows are the vectors being compared; we will refer to this organization as class-vector structure (**Figure 1a**). For example, consider the responses of different retinal neurons to the same set of visual stimuli. In this case, each visual stimulus can be considered a class (column) and each neuron (row) a vector of responses to the different classes (i.e., the set of stimuli). For any such dataset, PRED provides a unified measure of the similarity between the vectors and the separability of the classes. Put simply, class-vector PRED measures whether vector A’s value in a class is more similar to vector B’s value in the same class than to B’s value in another class. PRED is high when the distances are larger between values belonging to different vectors and different classes than between values belonging to different vectors but the same class (**Figure 1a**). In other words, a high value of PRED means that the two vectors have values not only with similar magnitudes but also with similar patterns across the classes. A zero value of PRED indicates that the two vectors have unrelated patterns across the classes. A negative value of PRED indicates that the two vectors have opposite patterns across the classes. Unlike correlation, PRED also accounts for the absolute differences between the values in the given vectors.

**Figure 1:**
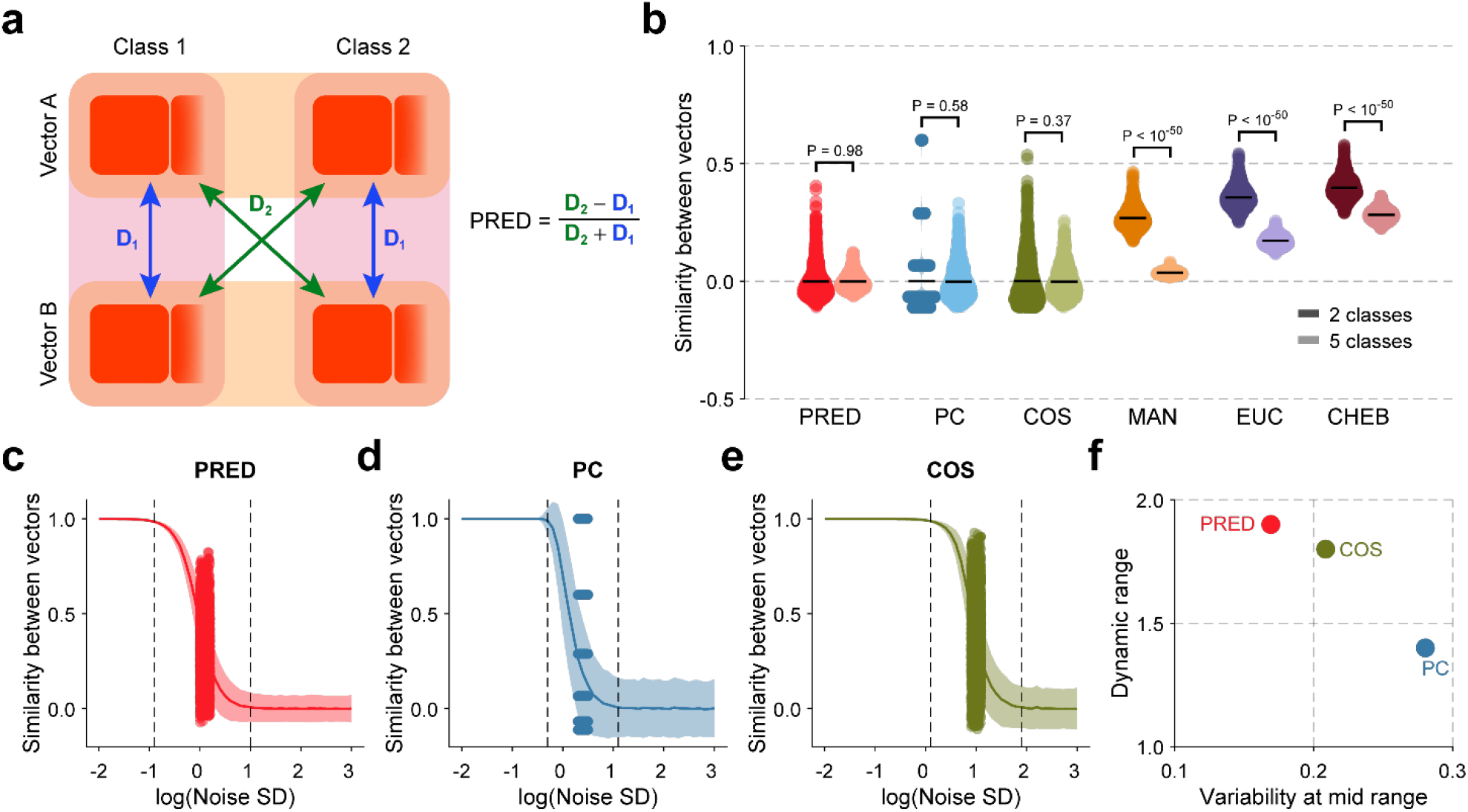
PRED is a robust metric for the assessment of similarity across vectors. **a** Schematic representation of Pairwise Relative Distance’s (PRED) calculation for a class-vector dataset. **b** Violin plots showing the chance level of each metric with simulated datasets containing 2 (darker colors) or 5 (lighter colors) classes. Each point within a violin represents the metric’s value for a different random seed (n = 1000 simulations for each number of classes). Note the change in the chance level of MAN, EUC, and CHEB metrics with the number of classes. PRED: Pairwise relative distance, PC: Pearson’s correlation, COS: Cosine similarity, MAN: Manhattan distance, EUC: Euclidean distance, CHEB: Chebyshev’s distance. Black horizontal line represents the mean. Error bsars represent s.e.m. **c—e** Change in the value of PRED **(c)**, PC **(d)**, and COS **(e)** with increasing noise level (shown on a log scale) in a simulated dataset with 2 classes and 10 individuals. The dark line shows the mean value over all simulations at the specified noise level (n = 1000 simulations per noise level). The shaded area represents 1 standard deviation around the mean. The two dashed vertical lines represent the boundaries of the dynamic range. Each point represents a different random simulation at the noise level corresponding to the mid-point of the dynamic range. **f** The dynamic range and the variability at the mid-point of the dynamic range are shown for each metric. PRED showed the highest dynamic range and the lowest variability.

We compared PRED and five other metrics on their ability to report the similarity across vectors within a class-vector dataset. These five metrics included Pearson’s correlation (PC), Cosine similarity (COS), Manhattan distance (MAN), Euclidean distance (EUC), and Chebyshev’s distance (CHEB). PRED, PC, and COS values range between −1 and 1, where 1 denotes high similarity; MAN, EUC, and CHEB range from 0 to ∞, where 0 denotes high similarity. To enable a direct comparison of the values of all these metrics, we transformed the distance-based metrics (MAN, EUC, and CHEB) to a range between 0 and 1 using a negative exponential (see **Materials and Methods**), such that 1 denotes high similarity for all the metrics **(Supplementary Figure 1a (i)**). We use the transformed distance-based metrics in all subsequent analyses unless otherwise stated.

For interpreting the values of a metric, it is helpful to know its chance level, i.e., the metric’s expected value for random data. For example, suppose a metric’s observed value for a given dataset is high relative to its chance level. In that case, one can reasonably infer that the vectors in the dataset have a high similarity: the more the difference, the higher the similarity. It is further desirable that the chance level remains unchanged with the size of the dataset (the number of classes in the dataset) so that values obtained from different datasets, regardless of their size, can be directly compared. To test each metric’s chance level, we simulated two different random datasets, one with 2 and the other with 5 classes. Each dataset included 10 vectors (with length equal to the number of classes) sampled from a uniform distribution between 0 and 1, ensuring no inherent similarity between vectors and difference between classes (see **Materials and Methods** for details). Expectedly, the observed chance level of PRED, PC, and COS was nearly 0 for both the 2-class and 5-class datasets; it was greater than 0 for MAN, EUC, and CHEB for both types of datasets (**Figure 1b**). Moreover, MAN, EUC, and CHEB’s chance levels were different for the datasets with different numbers of classes (**Figure 1b**). This difference occurs because the distances between vectors depend on the vectors’ sizes; we can more directly observe this change in chance levels with untransformed MAN, EUC, and CHEB metrics, all of which showed larger values with more classes (**Supplementary Figure 1b**). We tried to normalize these metrics according to the number of classes – for example, by dividing MAN by the number of classes or dividing EUC values by the square root of the number of classes. Although these normalizations reduced the overall differences between the chance levels for different numbers of classes, the differences remained significant (**Supplementary Figure 1c**). Thus, distance-based metrics do not provide a stable chance level.

Another important consideration for assessing a metric’s utility is its ability to report the level of similarity for a dataset, and its modifications, in a way that matches intuition. We had previously reported PRED’s advantages over PC in calculating stereotypy (Mittal et al., 2020). Here, we extend this analysis to include the other metrics. If the responses in a vector are the same for both classes, PRED reports a value of 0; however, PC is undefined, and COS reports a high value (**Supplementary Figure 1a (ii)**). If the two vectors exhibit opposite patterns across the classes (**Supplementary Figure 1a (iii)**), PRED and PC appropriately quantify the similarity as −1. COS, however, still reports a value close to 1, which does not match the intuitive difference between the two vectors. The distance-based metrics also fail to capture this difference: they report the same values of similarity in **Supplementary Figure 1a (iii)** and **(iv)**, even though in one case the vectors exhibit opposite patterns and in the other case they exhibit similar patterns across the two classes. If we linearly transform all the values in a dataset in the same manner, intuitively, the similarity between them should not change. Except for COS, all metrics are stable to global translational change, i.e., the addition of a constant to all the values in the dataset (**Supplementary Figure 1a (v)** compared to **(iv)**). Similarly, all metrics, except MAN, EUC, and CHEB, are stable to scaling modifications, i.e., multiplication of the entire dataset by a constant value (**Supplementary Figure 1a (vi)** compared to **(iv)**).

Overall, PRED behaved intuitively for various modifications within the datasets, while each of the other metrics deviates from the intuition in one or more cases (summarized in **Table 1**). As the distance-based metrics (MAN, EUC, and CHEB) lack a stable chance level, are not sensitive to patterns in the dataset, and are not robust to simple scaling transformations, we exclude them from further consideration as metrics of similarity.

**Table 1:** Summary of the properties of class-vector metrics.

| Metric | Range | Chance Level | Discreteness | Consistency with global scaling | Consistency with global translation |
| --- | --- | --- | --- | --- | --- |
| PRED | [-1 1] | 0 | Continuous | Constant | Constant |
| PC | [-1 1] | 0 | Discrete for 2 classes | Constant | Constant |
| COS | [-1 1] | 0 | Continuous | Constant | Changes |
| MAN | [0 1] | $> 0^a$ | Continuous | Changes | Constant |
| CHEB | [0 1] | $> 0^a$ | Continuous | Changes | Constant |
| EUC | [0 1] | $> 0^a$ | Continuous | Changes | Constant |

Values in red represent less desirable behavior compared with PRED.
<sup>a</sup> Chance level of these metrics varies with the number of classes.

Experimental datasets are often noisy. With any metric, we expect the similarity between two vectors to decrease as the noise level in the dataset increases, eventually reaching the chance level for extreme levels of noise. We studied how PRED, PC, and COS behaved for different noise levels using two parameters: dynamic range and variability. Here, dynamic range denotes the range of noise levels within which a metric exhibits unsaturated values and thus remains useful. Variability represents the sensitivity of a metric to noisy data: we consider a metric to have high variability if it shows very different values for different samples of the data at a given noise level. We quantified variability as the percent standard deviation over repeated simulations with noise at the mid-point of the dynamic range (see **Materials and Methods**). A useful metric should have a high dynamic range and low variability. We measured both these parameters for PRED, PC, and COS in a simulated dataset (see **Materials and Methods**) with increasing noise levels (**Figures 1c—e**). We found that PRED exhibited the highest dynamic range and lowest variability among all the metrics (**Figure 1f**). Even for simulated datasets with different base means, PRED was consistently more robust than the other metrics (**Supplementary Figures 1d, e**). Thus, PRED remains informative across a relatively large range of noise levels in the dataset and provides a relatively stable estimate of similarity.

### PRED for behavioral similarity assessment

We previously applied PRED to comparing the similarity of neural response patterns to an odor set across individuals (Mittal et al., 2020). However, in principle, it can be applied to any dataset where the data are arranged as vectors (each vector’s length equals the number of classes). Many behavioral studies examine if the behavioral outcomes of multiple individuals are similar over different time points. Here, one could consider the individuals as classes and each time point as a vector. Honegger et al. (Honegger et al., 2020) measured the preference indices of 141 *Drosophila* flies in a two-choice assay between two odors (3-octanol versus 4-methylcyclohexanol) over two different time points 24-hours apart (**Figure 2a**). They used PC to compare the similarity of preference index vectors across the two time points and found a moderate positive value of 0.35 (Honegger et al., 2020). Using PRED on the same data, we observed a value of 0.19, indicating a moderate similarity between behavioral preferences across the two time points.

**Figure 2:**
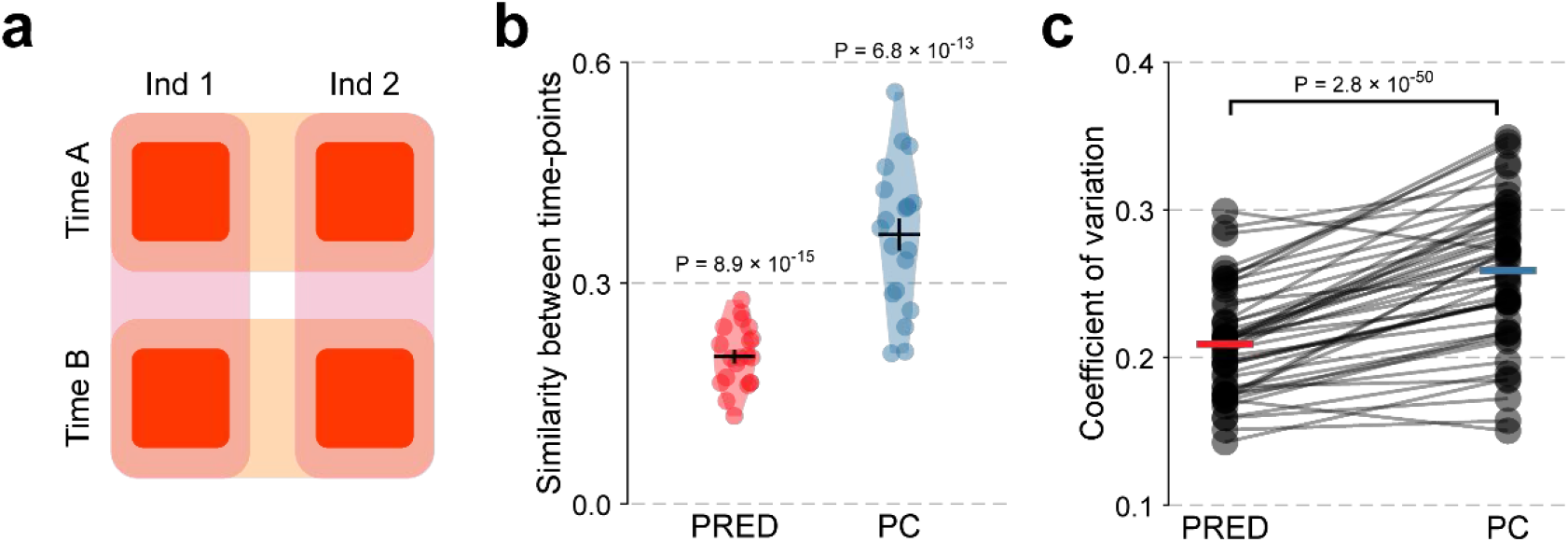
PRED is a suitable metric for measuring behavioral similarity. **a** Illustration of an individual-time dataset where each value represents the preference index of an individual animalat the specified time. **b** Across-time similarity in the MCH-OCT preference index of *Drosophila* measured with 70 individuals and 2 time-points. The 70 individuals were randomly sampled from a dataset with 141 individuals. The coefficient of variation (COV) is also displayed. Each point within a violin represents the mean similarity for a new randomly sampled dataset (n = 20 samplings). Black horizontal line represents the mean. **c** Coefficient of variations of 100 different repetitions of the analysis performed in **(b)**. Horizontal lines represent the mean COV over all repetitions (n = 50 repetitions). Lines connect the PRED and PC values from the same repetition.

Our results above (**Figure 1f**) have indicated that PRED is more stable than PC for noisy data. Therefore, we reasoned that it would also be more robust when working with incomplete datasets. The 141-fly behavioral dataset provided a suitable test case for this idea. We randomly selected 70 flies from the dataset and calculated the similarity of the preference index vectors at the two time points using PRED and PC. This random sampling was repeated 20 times, each resulting in a different value of PRED and PC. Even with incomplete datasets, both metrics reported significant similarity: 0.20 ± 0.04 (P = 8.9 × 10^−15^, n = 20; one sample t-test compared to 0) for PRED; and 0.37 ± 0.10 (P = 6.8 × 10^−13^, n = 20) for PC. Note that the PRED values were less variable (smaller s.d.) over the repeated samplings. Even the coefficient of variation, defined as 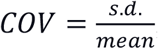, over these 20 samplings was smaller for PRED (0.21) than PC (0.27) (**Figure 2b**). Since these observed values of the COV may depend on the specific 20 samplings that occurred, we repeated the whole process of 20 samplings a total of 50 times and each time calculated the COVs for both metrics. This analysis confirmed that the COV was consistently lower for PRED (P = 2.8 × 10^−15^, n = 50, two-sample paired t-test; **Figure 2c**). Thus, PRED provides a relatively stable estimate of similarity for partial samplings of the dataset.

### Similarity in multi-dimensional data

So far, we have calculated similarity between two vectors where each vector contains a set of values corresponding to the set of classes—for example, comparing the response of a neuron to 2 stimuli (classes) in 2 individuals (vectors). This formatting is feasible for datasets where the response is a single number, such as the total number of spikes (or the net firing rate) evoked by a stimulus within a pre-defined time window. However, one may want to look at the response in finer detail, for example, by considering the temporal pattern of spikes evoked by the stimulus. We can represent the temporal pattern as a set of numbers by dividing the time window into, say, 10 bins and then counting the spikes in each bin. Thus, the response to a stimulus is now itself a 10-element vector rather than a single number (**Figure 3a**). In this case, if we want to compare the responses to a set of stimuli in two individuals, we need to compare two vectors of vectors rather than two vectors of numbers (**Figure 3a**).

**Figure 3:**
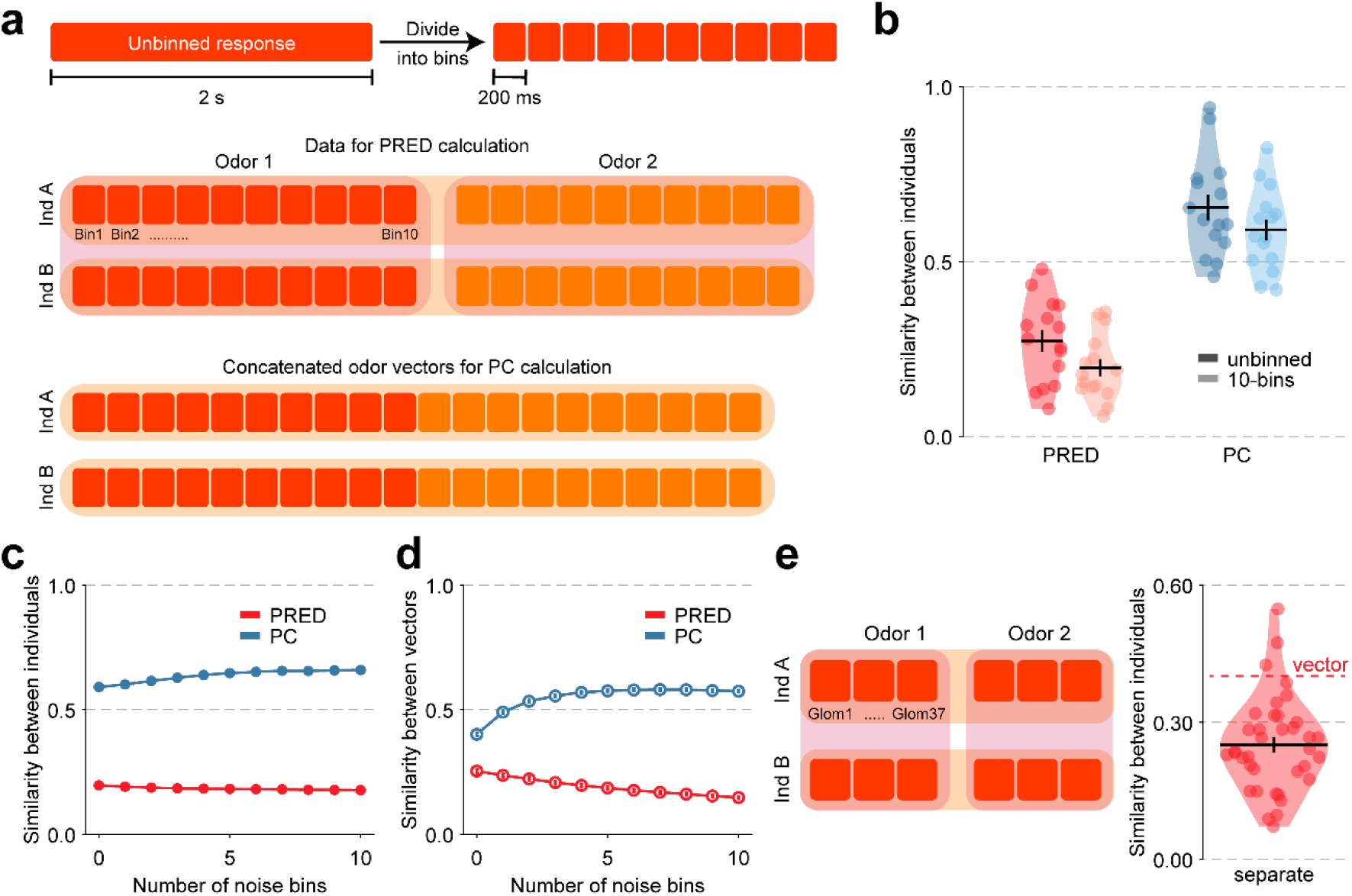
PRED natively supports multi-dimensional data. **a** Illustrations showing the unbinned and the 10-bin temporal vectors used for calculating the response similarity between individuals. For calculating PRED, the Euclidean distance between the 10-bin vectors across individuals is calculated. However, for calculating PC, the responses for both odors are first concatenated into a single 20-bin vector and then correlated across individuals. **b** Across-individual similarity when the neural response is quantified as a single unbinned number (darker colors) or as a 10-bin temporal vector (lighter colors). The data is taken from locust bLN1 neural responses (Gupta et al. 2014). Each point within the violin represents the similarity for a pair of individuals (n = 15). Black horizontal lines represent the mean, and error bars represent s.e.m. in all panels. **c** Across-individual similarity as a function of the number of extra bins (containing mostly noise) added to the original 10-bin vector for the same dataset as in **(b)**. Note that the similarity value reported by PC increases with the increasing number of bins. **d** Across-individual similarity as a function of the number of extra bins (containing noise) added to a 10-bin vector for simulated data with 2 odors and 10 individuals. The value in each extra bin is taken from a normal distribution with 0 mean and 1 s.d. Open circles denote the mean over 100 different random simulations. The similarity gradually reduces with the increasing number of noisy bins for PRED but increases for PC. **e** Illustration of the odor-individual dataset used for comparing the population response across individuals. Each bin represents the response of a glomerulus (Glom) in an individual for the odor tested. Violin plot shows the across-individual similarity measured by odor-individual (class-vector) PRED in a database with a population of 37 neurons, either considered separately (violin plot, where each point represents the PRED value for a neuron, n = 37) or considered together as a population vector (red dashed line).

Although correlation is frequently used to quantify the similarity between vectors, it is not equipped to handle vectors of vectors. A common modification to use correlation in such cases is concatenating the internal vectors within the outer vector to result in a single (and long) vector. In the example discussed earlier, it would mean combining the two 10-element vectors corresponding to the two stimuli to obtain a 20-element vector for each individual and then calculating the correlation between the 20-element vectors of the two individuals (**Figure 3a**). On the other hand, PRED is natively equipped to handle vectors of vectors and does not require concatenation: it involves calculating Euclidean distances between the values, which we can do irrespective of whether the values are single numbers or vectors. In the example discussed above, we can calculate *D*_1_ and *D*_2_ for PRED based on the 10dimensional Euclidean distances between the binned responses and then PRED using the regular formula, 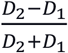 (**Figure 3a**).

We used both PRED and PC to compare the firing rates or the 10-bin temporal patterns evoked by odors in different individuals (see **Materials and Methods**). We performed this analysis in two different datasets: the olfactory response of mushroom body output neuron, bLN1, in locusts (Gupta and Stopfer, 2014) and four different projection neurons in *Drosophila* (Shimizu and Stopfer, 2017). We used a 2-second window after odor-onset to calculate the responses; in these datasets, the responses typically returned to baseline within 2 seconds in response to the 1-s odor pulse. Therefore, we can consider any spikes observed after this window as a part of the background spiking. For the temporal response, we divided this response into ten bins, each of length 200-ms (**Figure 3a**). Both PRED and PC revealed significant similarities between individuals and showed that the similarity was slightly lower when considering the temporal patterns instead of only the firing rates (**Figure 3b** and **Supplementary Figures 2a—d**).

Although PRED and PC behaved similarly in this analysis, PC can run into problems because of the concatenation step. Concatenation removes the distinction between the values belonging to different bins within the same class and the values belonging to different classes. For example, after concatenation, analyzing the 10-element temporal responses to 2 stimuli becomes identical to analyzing the firing rate responses to 20 independent stimuli, with each element contributing equally to the correlation. To illustrate why this can be problematic, we consider the case when the temporal response includes bins beyond the stimulus-evoked response; these bins would be mostly empty except for some noise. Since empty bins are similar by nature, including such bins in the response vectors and effectively treating them as independent stimuli after concatenation would spuriously increase the observed correlation.

In contrast, the calculation of Euclidean distances in PRED would be minimally affected by the empty bins: the distances would only become slightly noisier by the noise in the empty bins. Thus, PRED would report slightly lower similarity, which is a more intuitive outcome given the inclusion of irrelevant bins. To test these predictions in the actual datasets analyzed here, we included extra bins after the initial 10 bins of 200 ms duration. For example, in an 11-bin response, the first 10 bins would contain the first 2-s response after odor onset, while the last bin would contain an extra 200-ms response from 2 to 2.2-s after odor onset. Since the stimulus-evoked response typically lasted for less than 2 s, the extra bins included after the 2-s response are usually empty except for some noise. We found that, as predicted, the PC values increased as we added more and more extra bins in the response, whereas the PRED values decreased (**Figure 3c** and **Supplementary Figures 2f—i**). We further simulated a dataset containing two odors and ten individuals. The first 10 bins contained a simulated temporal response, and the subsequent bins contained random noise (see **Materials and Methods**). There was a noticeable increase in the PC values in these simulations with an increasing number of extra bins (**Figure 3d**). The effect became more pronounced when we added empty bins (i.e., bins with a value of 0) instead of bins with normally distributed noise. In this case, PRED values were constant as the empty bins did not affect the distances in PRED calculations (**Supplementary Figure 2e**). These results illustrate the pitfalls in using concatenated vectors in PC and suggest that PRED is a better alternative when working with multi-dimensional data.

Another type of multi-dimensional data is population-level data, i.e., the response of, say, 6 neurons from the same neural layer from two individuals responding to two stimuli. To analyze such a case, we can either calculate the similarity separately for each neuron and then take the average or directly consider the 6-element population response vector for each individual and odor. We used PRED to compare these two approaches, using a published dataset of calcium imaging responses of 37 antennal lobe glomeruli responding to 36 pure odors in 61 individuals (Badel et al., 2016). The similarity observed between individuals using the population vectors was significantly more than the average similarity of neurons considered separately (0.37 compared to 0.25 ± 0.10, P = 1.7 × 10^−10^, n = 37; one-sample t-test; **Figure 3e**). These results suggest that the combined cell population preserves more similarity within the system than individual cells, echoing previous studies’ results (Mittal et al., 2020). The results also illustrate the usefulness of PRED in analyzing population-level data.

### Class separability

The datasets we have considered so far had a class-vector structure (as shown in **Figure 1a**): multiple vectors (rows), each containing values for multiple classes (columns). The value of PRED for such a dataset depends on, and thus tells us about, both the similarity between the vectors and the separability of the classes. (Contrast this with Euclidean distance, which tells us only about the similarity between the vectors but is a poor indicator of class separability, as can be seen by comparing **Supplementary Figure 1a (iii)** and **(vi)**). In these datasets, there is a correspondence between the *i^th^* value in class 1 and the *i^th^* value in class 2, as they both belong to the same vector in row *i* (which could be an individual, a time-point, or any other variable depending on the experimental context). However, many datasets do not have this correspondence (i.e., there are no row-vectors) — for example, in neuroscience, one often measures the responses of a neuron or a brain region to different stimuli (classes) and takes multiple measurements (called trials or samples) for each stimulus. In such cases we are left with only classes (columns), with each class containing multiple values (as shown in **Figure 4a**). This formatting is commonly used in datasets with repeat measurements over multiple classes. Here, the numbers of samples for different classes do not have to be identical. Each sample value within a class may be a single number (e.g., the firing rate of a neuron or the preference index of an animal) or a set of numbers (e.g., a binned temporal response or a population response). Assuming that the samples within a class are generated under identical experimental conditions and that the samples in different classes are generated independently, there is no logical correspondence between the *i^th^* sample in class 1 and the *i^th^* sample in class 2. We will refer to such datasets as class-sample datasets. In such datasets, one often wants to know about the separability of the classes.

**Figure 4:**
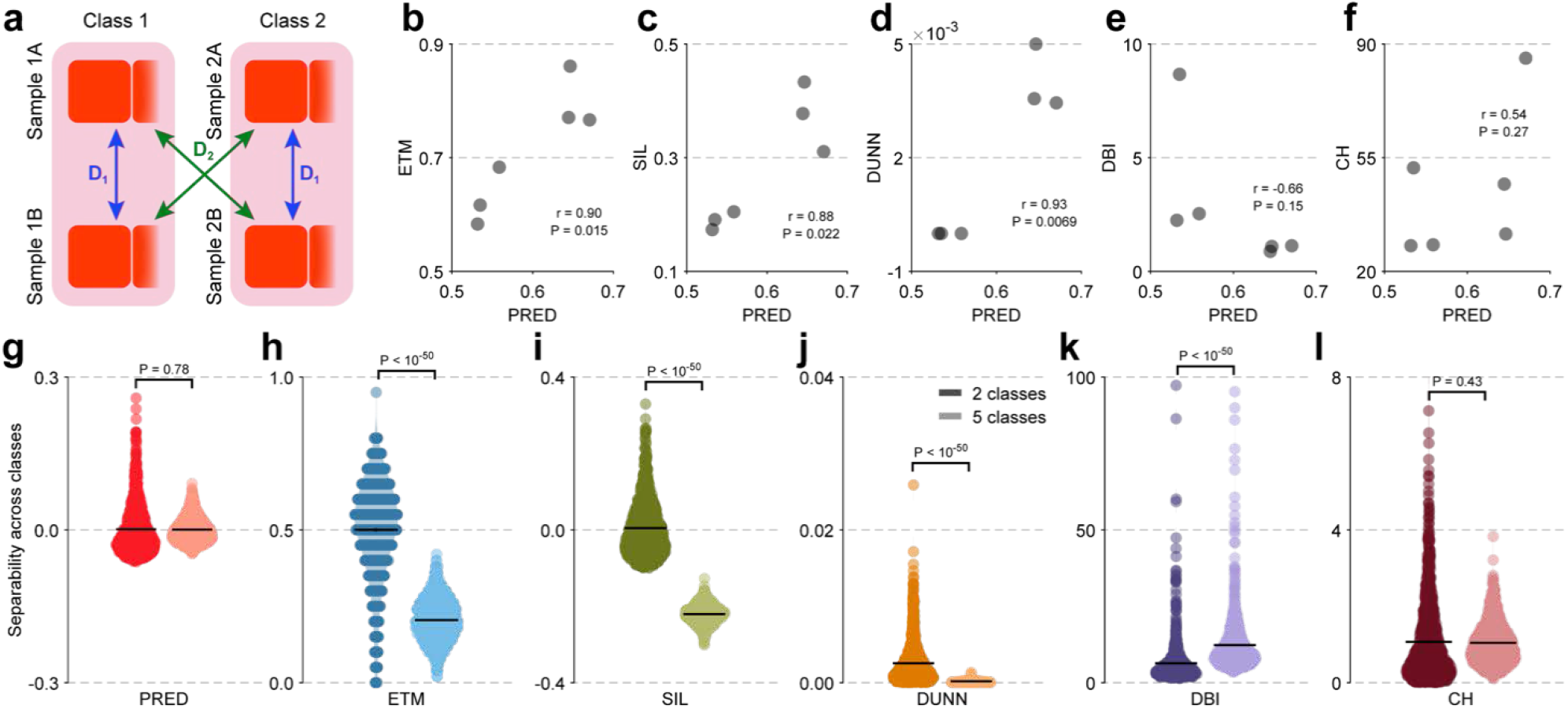
PRED is suitable for assessing class separability in class-sample datasets. **a** Schematic representation of Pairwise Relative Distance (PRED) calculation for a class-sample dataset. **b—f** Odor separability measured using PRED compared to that measured using other commonly used metrics. Each point corresponds to one individual in the dataset taken from locust bLN1 neural responses (Gupta and Stopfer, 2014) (n = 6 individuals). Note that PRED values were positively correlated with the values obtained from other metrics (DBI expectedly showed a negative correlation as DBI’s polarity is inverted). ETM: Euclidean template matching, SIL: Silhouette index, DUNN: Dunn’s index, DBI: Davies-Bouldin index, CH: Calinski-Harabasz index. **g—l** Violin plots showing the chance level of each metric with simulated datasets containing 2 (darker colors) or 5 (lighter colors) classes. Each point within a violin represents the metric’s value for a different random seed (n = 1000 simulations for each number of classes). Note the change in the chance level of all metrics except PRED and CH with the number of classes. Black horizontal line represents the mean. Error bars represent s.e.m.

A similar requirement arises when evaluating the output of unsupervised clustering algorithms, which use statistical methods to divide a collection of values into different clusters. The resulting clusters are analogous to classes in the above formulation, and their assigned members are analogous to samples. Here also, one often wants to know how well separated the observed clusters are. For example, Karagiannis et al. classified neuropeptide Y-expressing neocortical interneurons into 3 different types based on their morphology using a K-means clustering algorithm (Karagiannis et al., 2009). They then used the Silhouette index (Rousseeuw, 1987) to evaluate the quality of the clustering obtained. Another study used the Silhouette index to assess the efficiency of single nucleotide polymorphism genotyping assays in dividing samples into 3 different groups: homozygous for the first allele, homozygous for the second allele, or heterozygous (Lovmar et al., 2005). Apart from the Silhouette index (Rousseeuw, 1987), an evaluation of a clustering technique’s efficacy can be made using other internal clustering validation indices like the Davies-Bouldin index (Davies and Bouldin, 1979) or the Dunn’s index (Dunn, 1974). Another method commonly used to measure class separability is Euclidean template matching (ETM), which involves classifying each value based on its Euclidean distance from class templates (constructed from the remaining data) and then calculating the average accuracy from these classifications (Stopfer et al., 2003).

Since the PRED value for a class-vector dataset depends on class separability, we asked whether PRED can also be used as a measure of class separability in class-sample datasets (**Figure 4a**). We compared PRED to five commonly used metrics: Silhouette index (SIL), Davies-Bouldin index (DBI), Dunn’s index (DUNN), ETM, and Calinski-Harabasz index (CH) (see **Materials and Methods** for a description of each metric). As an initial test of PRED’s feasibility for this application, we used two different datasets containing repeated responses to different odors. We obtained one dataset from the identified *bLN1* neuron in locusts (Gupta and Stopfer, 2014) and another from four identified projection neurons in *Drosophila* (Shimizu and Stopfer, 2017). Each dataset contains the response from multiple individuals; we compared the odor separability calculated using PRED and the other metrics for each individual. We found that PRED values were somewhat correlated with the values from other metrics in both the datasets (**Figures 4b—f** and **Supplementary Figures 3a—e**). (Note that the correlation with DBI is negative because a lower DBI value indicates a higher separability, whereas the opposite is true for PRED and the other four metrics). These correlations with the established metrics suggested that PRED might also be useful as a metric of class separability. To explore this further, we compared PRED’s performance with the other metrics in various situations.

As discussed in the analysis of class-vector datasets, a key feature of any metric is its chance level. For evaluating the chance level of separability metrics in class-sample datasets, we simulated datasets containing clusters (classes) of points with fixed radii on a 2-d plane and different levels of noise (**Supplementary Figure 3f**; see **Materials and Methods** for details). As we increase the noise in the simulated dataset, the classes lose their separability (**Supplementary Figures 3f—h**). We used datasets with extremely high noise levels to calculate the chance level of each of the six metrics. Further, we checked how the chance levels depend on the number of classes in the dataset. PRED showed a chance level close to 0, regardless of the number of classes. CH showed a chance level greater than 0 that was not different for 2-class or 5-class datasets (**Figures 4g, l**). However, the chance levels of the other four metrics changed significantly with the number of classes (**Figures 4h—k**).

Imagine a large dataset containing many classes where any two classes have the same level of separability, whose value is not known to us. Further, imagine that, for practical reasons, we have access to only a subset of the dataset covering some of the classes, and our task is to use different metrics to estimate the class separability. An ideal metric should estimate the same underlying class separability, regardless of the number of classes available in our subset. To check how the six metrics under consideration perform on this criterion, we simulated a dataset with a low level of noise (thus with reasonable class separability) and varied the number of classes. We found that the separability reported by all metrics except PRED varied with the number of classes (**Figure 5a**).

**Figure 5:**
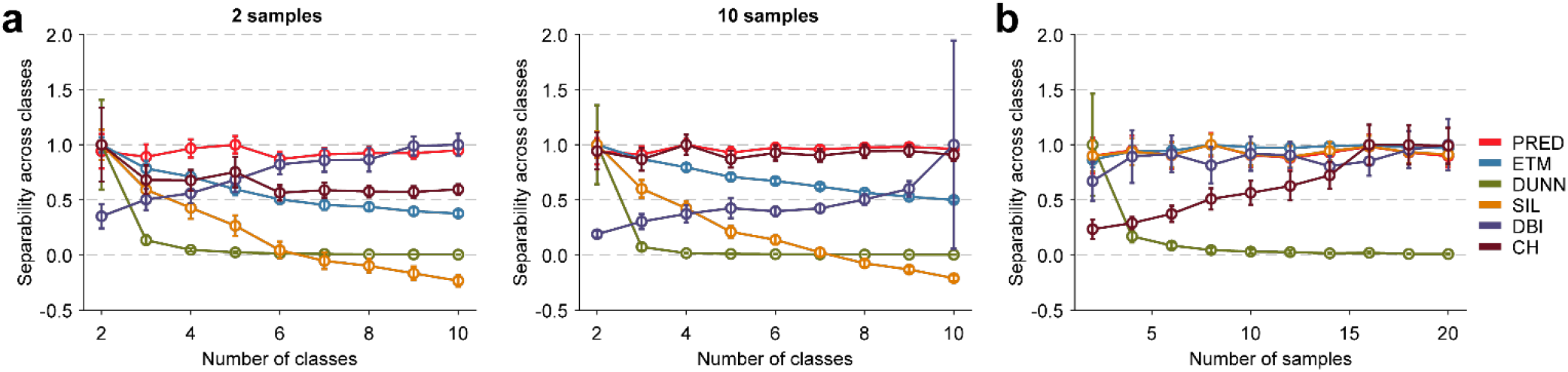
Unlike PRED, most other metrics vary with an increasing number of classes or samples. **a** Class separability as a function of the number of classes using simulated data with 2 samples (left) or 10 samples (right). Each metric was normalized by its maximum value observed among the mean values for different numbers of classes. Note that all metrics except PRED and SIL show change with the increasing number of classes. Open circles denote the mean value over 100 different random simulations for the specified numbers of classes, and error bars denote s.e.m. **b** Similar plot as in (**a)** but with 2 classes and an increasing number of samples (n = 100 simulations for each number of samples). Note the change in the value of ETM, DUNN, and CH with an increase in the number of samples. Also, in all plots, DBI values show an opposite trend as compared with the other metrics because DBI is higher for less separable classes.

CH values decreased with the number of classes when we had 2 samples per class but not when we had 10 samples per class (**Figure 5a**; **Figure 4l** also had 10 samples per class, which explains no change in the CH chance level). This result indicated that the number of samples could also bias the value of a metric. Ideally, the separability of the classes should not depend on how many samples are available for each class. For example, our estimate of how well a neuron can differentiate two sensory stimuli (a property of the neuron and the stimuli) should not be biased by the number of recording trials available (an experimental parameter). We performed another set of simulations with 2 classes and an increasing number of samples per class. We found that CH, ETM, and DUNN values varied significantly with the number of samples (**Figure 5b**), while PRED, SIL, and DBI were relatively stable. We conclude that PRED provides an unbiased estimate of class separability regardless of the number of classes or the number of samples per class. Therefore, we can reliably use it with datasets of all sizes.

We next studied the stability of each metric against noisy data by checking the dynamic range and the variability at the midpoint of the dynamic range. We simulated datasets with noise levels ranging from zero (highly separable classes) to very high (poorly separable classes). As before, we estimated the dynamic range as the range of noise levels for which a metric remained unsaturated and variability as the percent standard deviation over repeated simulations with noise at the mid-point of the dynamic range (**Figures 6a—f**). PRED and SIL showed the best combination of large dynamic range and small variability (**Figure 6g**). DUNN had the lowest dynamic range and high variability, while DBI exhibited a high dynamic range but also the highest variability (**Figure 6g**). We used the Drosophila and locust datasets to complement the simulation results. We added increasing amounts of noise to each value in the datasets and then compared the metrics (**Supplementary Figure 4**; see **Materials and Methods**). Again, PRED and SIL exhibited large dynamic ranges and small variabilities in all cases. DUNN and DBI showed a high dynamic range in some cases but were the worst performers in variability in most neurons. Overall, PRED and SIL appear to be the most robust metrics in handling noisy datasets. Considering that SIL values (including the chance level) depend on the number of classes, as discussed above, PRED appears to be the best among the considered metrics for quantifying class separability (summarized in **Table 2**).

**Figure 6:**
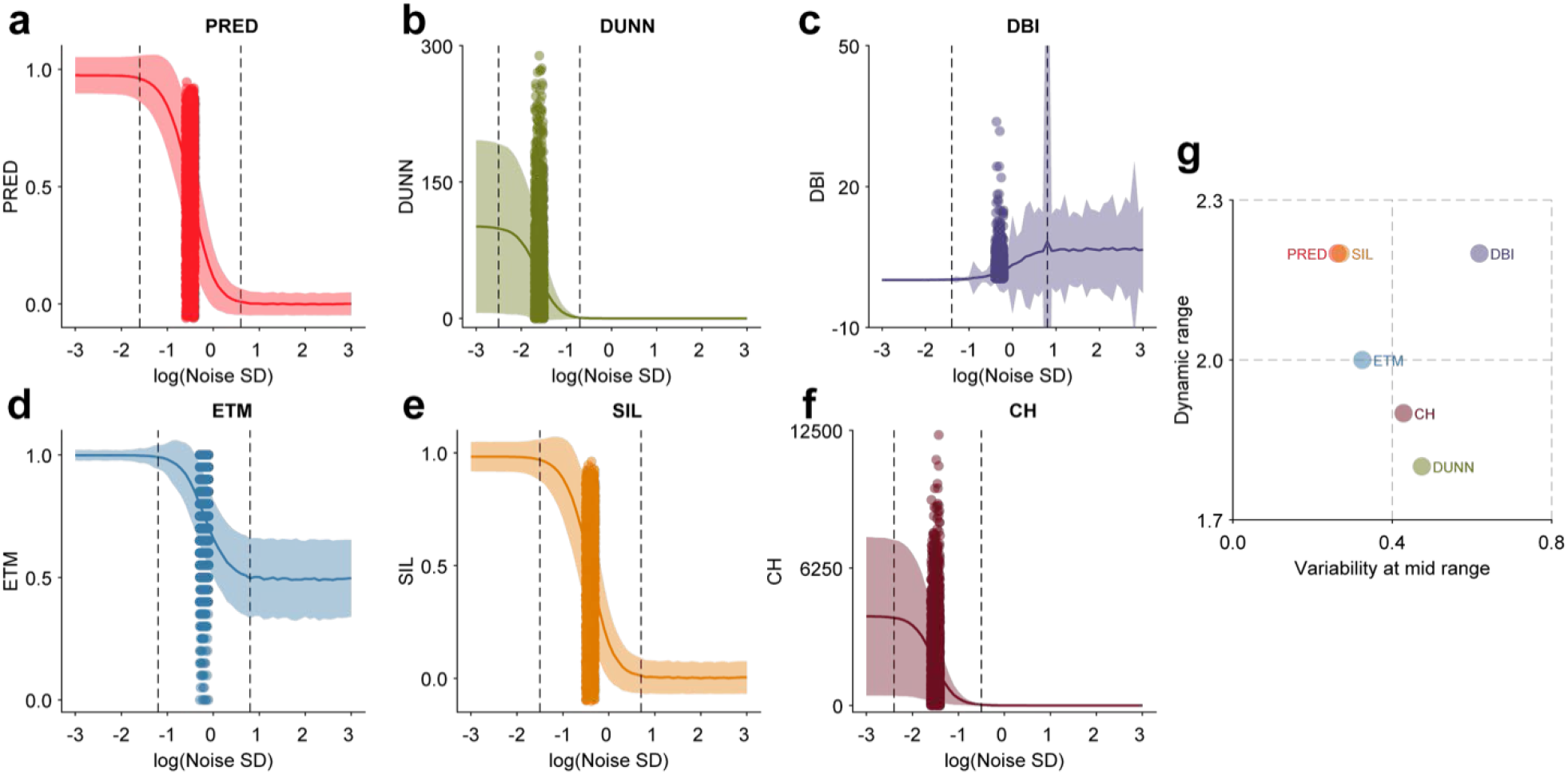
Comparison of dynamic range and variability of class-sample metrics. **a—f** Change in the value of PRED **(a)**, DUNN **(b)**, DBI **(c)**, ETM **(d)**, SIL **(e)**, and CH **(f)** with increasing level of noise (shown on a log scale) in a simulated dataset with 3 classes and 10 samples. The solid trace shows the mean values over all simulations for each noise level (n = 1000 simulations per noise level). The shaded area represents 1 s.d. around the mean. The dashed vertical lines represent the boundaries of the dynamic range. Each point represents a different random simulation at the noise level corresponding to the mid-point of the dynamic range. **g** The dynamic range and the variability at the mid-point of the dynamic range are shown for each metric. PRED showed a reasonably large dynamic range and low variability.

**Table 2:** Summary of the properties of class-sample metrics.

| Metric | Range | Chance Level<sup>a</sup> | Discreteness | Consistency with number of classes | Consistency with number of samples | Dynamic range<sup>b</sup> | Variability<sup>b</sup> |
| --- | --- | --- | --- | --- | --- | --- | --- |
| PRED | [-1 1] | 0 | Continuous | Constant | Constant | - | - |
| ETM | [0 1] | 0.5<sup>c</sup> | Discrete | Changes | Changes | Smaller | Higher |
| SIL | [-1 1] | 0<sup>c</sup> | Continuous | Changes | Constant | Similar | Similar |
| DBI | [Inf 0] | >0<sup>c</sup> | Continuous | Changes | Constant | Smaller | Higher |
| DUNN | [0 Inf] | >0<sup>c</sup> | Continuous | Changes | Changes | Smaller | Higher |
| CH | [0 Inf] | >0<sup>c</sup> | Continuous | Changes | Changes | Smaller | Higher |

Values in red represent less desirable behavior compared with PRED.
<sup>a</sup> reported for a dataset with 2 classes
<sup>b</sup> as compared to PRED
<sup>c</sup> Chance level of these metrics varies with the number of classes.

Class separability depends on how different the values are across the classes and how similar they are for different samples within each class. PRED, thus, may be a useful metric when both within-class similarity and across-class differences are analyzed simultaneously. Kermen et al. (Kermen et al., 2020) looked at zebrafish olfactory behaviors elicited by a set of 18 odors in different individuals while performing 4 repeated trials with each odor. They calculated the intra-individual similarity by correlating the behavioral responses across all pairs of trials for each individual and the inter-individual similarity by correlating the trial averaged response of all pairs of individuals. Then they looked at pairs of these two similarity values to examine how consistent the responses produced by each odor were within and across individuals. If one wants to know which odors produce relatively similar responses within individuals but different across individuals, PRED can provide the answer with a single number. We calculated PRED considering individuals as classes and trials as samples (**Figure 7a**; see **Materials and Methods**). We found that the behavioral responses were relatively different across individuals and consistent across trials for these odors: cadaverine (0.39±0.34, P = 1.8 × 10^−6^, n = 28), blood (0.39±0.35, P = 8.2 × 10^−4^, n = 15), skin (0.26±0.32, P = 0.007, n = 15), bile (0.17±0.26, P = 4.2 × 10^−4^, n = 36), sperm (0.15±0.35, P = 0.007, n = 45), cysteine (0.13±0.38, P = 0.05, n = 36), and arginine (0.12±0.26, P = 0.01, n = 36) (**Figure 7b**).

**Figure 7:**
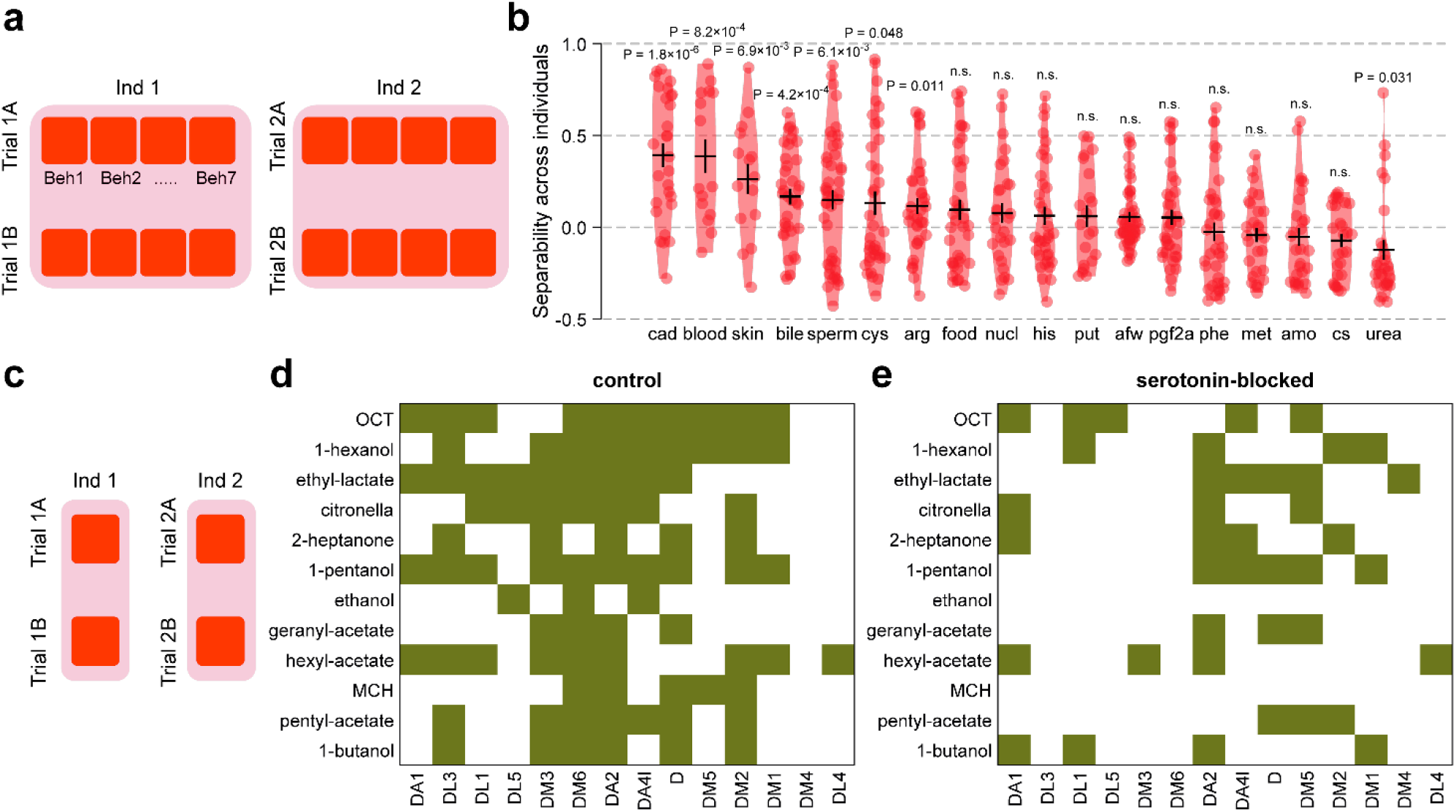
Using PRED to measure individuality of neural responses. **a** Illustrations of an individual-trial dataset where values in each column represent the repeated behavioral responses of an individual (to a particular odor). Each behavioral response is a 7-length vector, with each bin in this vector representing a specific physical behavior (Beh). **b** Individual-trial (class-sample) PRED for zebrafish behavioral data calculated separately for each odor. The odors are sorted from left to right in decreasing order of PRED value. Each point in the violin represents an individual pair (cad: n = 28, blood: n = 15, skin: n = 15, bile: n = 36, sperm: n = 45, cys: n = 36, arg: n = 36, food: n = 36, nucl: n = 28, his: n = 36, put: n = 21, afw: n = 45, pgf2a: n = 36, phe: n = 36, met: n = 28, amo: n = 28, cs: n = 28, urea: n = 28). Black horizontal line represents the mean. Error bars represent s.e.m. n.s. means not significant. **c** Illustration of an individual-trial dataset where values in each column represent the repeated responses of an individual (in a particular glomerulus and to a particular odor). **d, e** Individual-trial (class-sample) PRED for different PN-odor responses in control **(d)** and serotonin-blocked **(e)** *Drosophila*. Green color indicates PRED values significantly greater than 0, indicating good separability across individuals. Note the fewer number of green values after serotonin-blockage. Significance was measured using one-sample t-test.

### Using PRED for assessing individuality

Honegger et al. (Honegger et al., 2020) observed that odor preferences of *Drosophila* varied more across individuals than across trials within an individual. Consistent with this, they also found that the odor responses of the projection neurons were also more variable across individuals than across trials, suggesting that this response individuality may underlie the behavioral individuality. The behavioral individuality depended on serotonin: it reduced when the flies were fed alpha-methyl tryptophan, a serotonin synthesis blocker. However, somewhat unexpectedly, they did not detect a reduction in the response individuality in the presence of the serotonin blocker. Their analysis used principal component analysis and Bayesian modeling to compute inter-fly and intra-fly distances. Since quantifying individuality requires an assessment of inter-individual differences relative to intra-individual differences, we reasoned that individuality could be aptly described by class separability, where the individuals are classes, and the trials are samples within each class. We reanalyzed their data using individual-trial (class-sample) PRED to quantify the individuality of the PN responses to different odors (**Figure 7c**; see **Materials and Methods**). In the wild-type flies, we observed that 50% (84 out of 168) of the PN-odor responses were significantly separable across individuals (**Figure 7d**), matching the conclusions of Honegger et al. However, in serotonin-blocked flies, this fraction reduced to only ~24% (40 out of 168; **Figure 7e**) even though the original analysis was not able to uncover this reduction. Thus, our reanalysis of response individuality shows that serotonin indeed affects the PN response individuality. By resolving the contradiction between the behavioral data and the PN response data in the presence of serotonin blockage, our analysis using PRED lends additional support to the idea of Honegger et al. (Honegger et al., 2020) that PN response individuality determines behavioral individuality.

### Using PRED for analyzing connectomic data

Recent advances in high-throughput electron microscopy and image segmentation methods have made it possible to reconstruct neuronal morphologies and connections in large brain areas. For *Drosophila*, two public datasets, namely the full adult fly brain or FAFB (Zheng et al., 2018) and the Hemibrain (Scheffer et al., 2020), have recently become available. As these datasets are generated from two different individuals, they provide an opportunity for measuring stereotypy in the connectivity patterns of neurons across individuals. A recent study by Schlegel et al. (Schlegel et al., 2021) used these two datasets to measure stereotypy in the input connections received by the lateral horn neurons (LHNs) from the projection neurons (PNs). For each LHN, they calculated a vector of connectivity with different types of PNs and used the cosine metric (COS) to estimate the similarity between such vectors. They demonstrated stereotypy in the inputs of LHNs by a combination of two results: (i) when comparing LHNs belonging to the same cell type, the COS values for LHNs across the two datasets were high and similar to the COS values for LHNs within a dataset; and (ii) when comparing LHNs belonging to different cell-types, the COS values for LHNs across the two datasets were low and similar to the COS values for LHNs within a dataset.

PRED allows one-shot quantification of stereotypy in this case with a single number. Based on their morphologies and connections to other neurons, the LHNs have been grouped into ‘connectivity types,’ which are further grouped into ‘regions,’ ‘tracts,’ and ‘cell types’ in the increasing order of hierarchy (see **Materials and Methods**). Although it has not been possible to match the neurons in the two datasets unambiguously, these higher-order groupings have been labeled in both datasets. We computed a 57-length glomerular input vector for each group by averaging the connectivity vectors of all LHNs belonging to the group (**Figure 8a**). To estimate stereotypy in the glomerular input vectors of groups at a particular hierarchy level, we calculated the group-dataset (class-vector) PRED (**Figure 8a**). At the level of ‘connectivity types,’ we found that the PRED value was 0.56±0.25 (P = 4.4 × 10^−193^, n = 496), notably higher than the chance level of 0, suggesting that the averaged connectivity vectors were separable across connectivity types and similar across the two datasets. Similarly, high PRED values were also seen at other grouping levels (cell type: 0.56±0.25, P = 1.4 × 10^−147^, n = 378; tract: 0.61±0.16, P = 1.4 × 10^−40^, n = 66; region: 0.56±0.18, P = 5.5 × 10^−4^, n = 6), confirming the stereotypy in the connectivity patterns of LHNs groups across the two databases.

**Figure 8:**
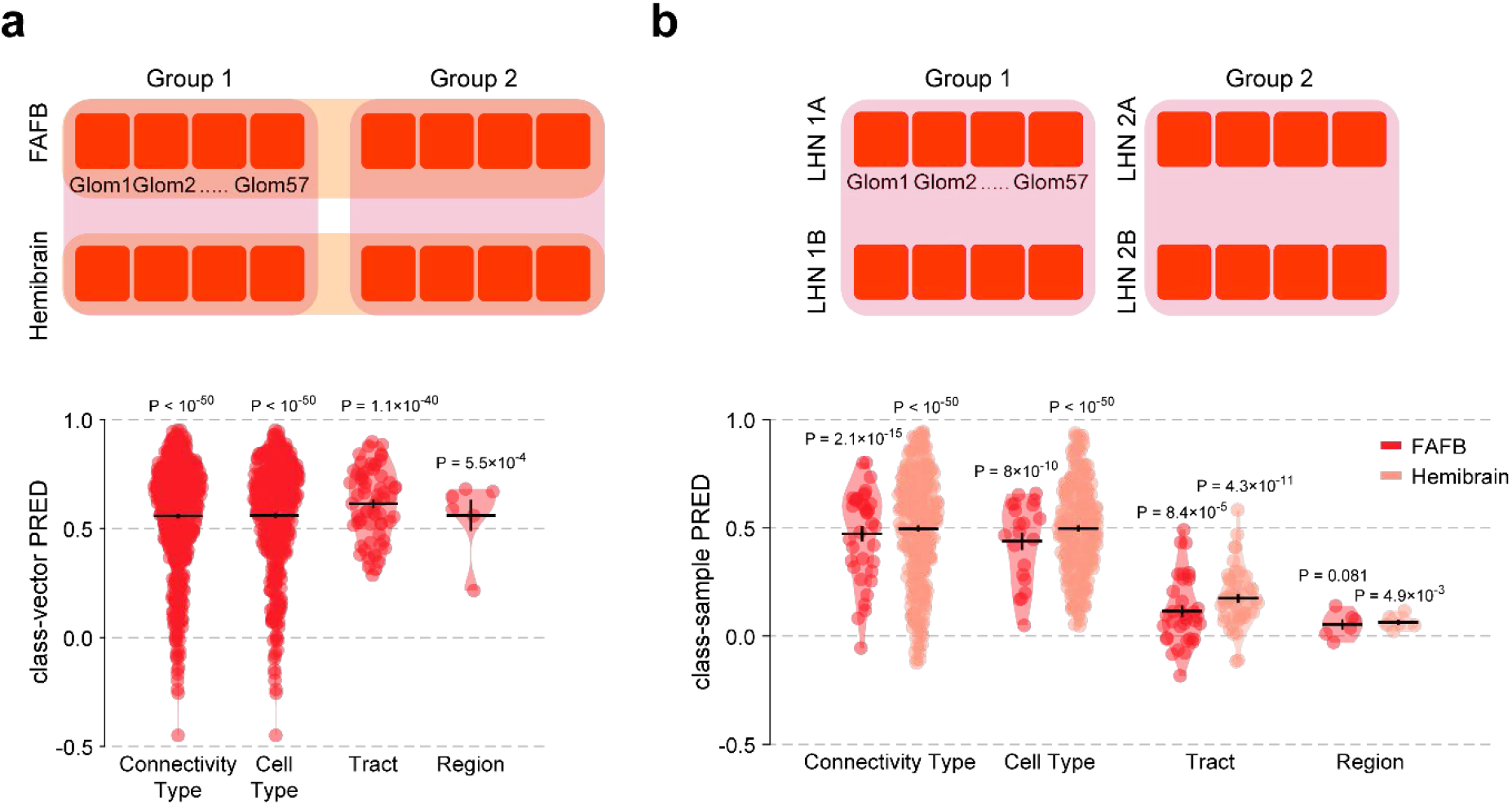
Using PRED as a measure of similarity and separability for connectomic data. **a** Illustration of the group-database (class-vector) structure used for comparing the two datasets, FAFB and Hemibrain. Each bin represents the average strength of connections between the LHNs belonging to the group and a single glomerulus (Glom). High value of group-database PRED confirms stereotypy between FAFB and Hemibrain datasets for all 4 levels of groupings of lateral horn neurons (LHNs). Each value in the violin represents a pair of groups within the specified hierarchy level (connectivity type: n = 496 pairs of connectivity types, cell type: n = 378 pairs of cell types, tract: n = 66 pairs of tracts, region: n = 6 pairs of regions). The calculations were performed over the antennal lobe glomerulus to LHN connectivity data. The connectivity values were averaged over all all neurons within the specified groups. **b** Illustration of the group-neuron (class-sample) dataset for calculating the across-group separability of neuron connectivity patterns. Each column contains the connectivity vectors of all LHNs belonging to a group. Each bin represents the strength of connections between an LHN and a single glomerulus. Group-neuron PRED for the dataset with individual neurons grouped into connectivity types (FAFB: n = 36 pairs of connectivity types, Hemibrain: n = 276), cell types (FAFB: n = 21 pairs of cell types, Hemibrain: n = 210), tracts (FAFB: n = 36 pairs of regions, Hemibrain: n = 45) or regions (FAFB: n = 6 pairs of tracts, Hemibrain: n = 6) for each of the two datasets, FAFB and Hemibrain. Black horizontal line represents the mean, and error bars represent s.e.m.

The above analysis compared the averaged glomerular connectivity patterns of different groups. Next, we sought to assess whether the glomerular connectivity patterns of different neurons within a group were more consistent than the patterns of neurons across different groups at the same hierarchy level. This could be easily quantified as group-separability using group-neuron (class-sample) PRED. In both the datasets, we found that the ‘connectivity types’ were highly separable (FAFB: PRED = 0.47±0.21, P = 2.1 × 10^−15^, n = 36; Hemibrain: PRED = 0.50±0.25, P = 3.7 × 10^−96^, n = 276; **Figure 8b**). Similarly, the cell types were also highly separable (FAFB: PRED = 0.44±0.19, P = 8 × 10^−10^, n = 21; Hemibrain: PRED = 0.50±0.21, P = 6.5 × 10^−88^, n = 210). The separability reduced as we went to higher levels in the group hierarchy, namely the ‘tracts’ (FAFB: PRED = 0.11±0.15, P = 8.4 × 10^−5^, n = 36; Hemibrain: PRED = 0.18±0.14, P = 4.3 × 10^−11^, n = 45) and the ‘regions’ (FAFB: PRED = 0.05±0.06, P = 0.081, n = 6; Hemibrain: PRED = 0.06±0.3, P = 0.0049, n = 6). This reduction in class separability reflects the increasing diversity of neurons within the higher-level groups. Overall, these results demonstrate how class-sample PRED can be used as a sensitive and easy-to-use metric of class separability.

## Discussion

Overall, we found that Pairwise Relative Distance (PRED) is a robust metric for quantifying vector similarity and class separability in class-vector datasets and offers several advantages over distance-based metrics, Pearson’s correlation, or cosine similarity. Importantly, PRED quantified the similarity in a consistent way close to our intuitive understanding of the data. Datasets in different studies often vary in terms of their size and the scale of the responses. If the similarity metric is affected by these parameters, it becomes difficult to compare the results obtained across studies. PRED, however, remained agnostic to the size of the dataset and was unchanged with global modifications of the data (**Figure 1** and **Supplementary Figure 1**). We can, thus, directly compare PRED values obtained from different studies.

Experimental studies may be limited in the amount of data that they can collect; in terms of, for example, how many different stimuli one can present, or how many individuals can study, or how many trials one could perform, and so on. Also, experimental data is subject to noise from multiple sources. Thus, it is desirable to analyze datasets with a metric that is robust to noise. In our study, PRED exhibited the largest dynamic range and the lowest variability among the metrics tested. It also worked well with incomplete datasets (**Figures 1, 2**, and **Supplementary Figure 1**).

Many metrics are available for calculating the similarity of vectors when each value within the vector is a scalar quantity (a number). However, we cannot directly use these metrics when each value within the vector is itself a vector (a set of numbers), as is the case with temporally patterned neural responses or population responses. One could forcibly convert the vector of vectors into a long vector of numbers through concatenation. However, concatenated vectors lose the distinction between classes and the elements of values within a class. As we showed by simulating increasingly longer temporal patterns, this can lead to an inaccurate estimation of similarity. On the other hand, PRED provides a more straightforward and intuitive method for analyzing multi-dimensional data while preserving the inherent relations between different dimensions (**Figure 3** and **Supplementary Figure 2**).

We found that PRED also works well for analyzing class separability in class-sample datasets, as the results with PRED were well correlated with those obtained from other commonly used metrics. PRED provided a stable chance level and was unaffected by the dataset’s size, whereas most of the other metrics that we tested varied with an increase in the number of classes or samples. We tested the robustness of several internal clustering validation metrics to noisy datasets. In these analyses using simulated and experimental data, PRED was consistently among the metrics with the highest dynamic range and the lowest variability. Thus, PRED presents a consistent and more reliable alternative for evaluating class separability in class-sample datasets (**Figures 4 – 8 and Supplementary Figures 3— 5**).

When dealing with large datasets, one consideration in choosing a metric is its computational time complexity. Since PRED calculates the similarity iteratively for all combinations of pairs of classes and pairs of vectors, its time complexity is of the order of 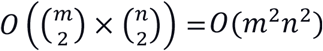, where *m* and *n* are the numbers of classes and vectors, respectively. Thus, the time required to compute PRED increases polynomially with an increase in the dataset’s size. Other class-vector metrics including Pearson’s correlation, cosine similarity, and distancebased metrics have *O*(*mn*^2^) time complexity. However, datasets in many applications are small enough (*m, n* ≤ 100) that the time complexity of PRED would not become a limiting consideration.

We originally designed PRED for class-vector datasets, in which there is a correspondence between the *i^th^* element in class 1 and the *i^th^* element in class 2, as both elements belong to the same vector (row). PRED calculation makes use of this correspondence when making the 2×2 matrices for a pair of classes: if a 2×2 matrix has the *i^th^* and the *j^th^* values from class 1, it must have the *i^th^* and the *j^th^* values from class 2). In class-sample datasets, this correspondence across classes is absent, as there is no ordering among the class elements – all samples are random replicates. This lack of order poses a dilemma while calculating PRED: which pair of values in class 2 should we use for making the 2×2 matrix with a particular pair of values in class 1? We overcome this dilemma by considering all possible pairs from class 2 iteratively for a given pair of values in class 1. This method (‘exhaustive PRED’) increases the time complexity from 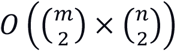 to 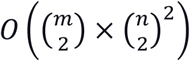 for class-sample datasets, assuming each of the *m* classes has *O*(*n*) elements (**Supplementary Figure 5a**). In practice, the extra time required for ‘exhaustive PRED’ would be noticeable only for large datasets with hundreds of classes and samples. The calculation can be made faster using an approximation (‘fast PRED’). In ‘fast PRED,’ we assign an arbitrary order to the elements in each class (e.g., the order in which the values were saved) and then create 2×2 matrices in the same way as is done in class-vector datasets: when we take the *i^th^* and the *j^th^* values from class 1, we also take the *i^th^* and the *j^th^* values from class 2. Using simulations (see **Materials and Methods**), we found that the difference between the ‘exhaustive PRED’ and the ‘fast PRED’ values was ~3% for datasets with more than 15 samples (**Supplementary Figure 5b**). Changing the ordering of elements within classes did not have a noticeable effect on the value of PRED. Thus, we can efficiently and reliably compute PRED for large class-sample datasets.

Class-sample PRED essentially compares the within and across class variation of samples. As classification is a very commonly used operation, there has been a strong interest in comparing various metrics under different scenarios (Arbelaitz et al., 2013; Brun et al., 2007; Guerra et al., 2012; Gurrutxaga et al., 2011; Niemelä et al., 2018). Apart from the metrics that we have already compared with PRED, other metrics with similar approaches, like the t-statistic or Fisher discriminant, can potentially be used for analyzing class-sample datasets. However, these metrics have their drawbacks. The calculation and the interpretation of the t-statistic depend on the degree of freedom, which is a function of the number of samples observed. The discriminant analysis assumes a linear separation between the classes and thus might not be ideal for neural datasets. Another approach, formulated by Huerta et al. (Huerta et al., 2004), also quantifies intra-class and inter-class differences. They calculated average within-class (*D_intra_*) and across-class (*D_inter_*) distances, similar to our *D*_1_ and *D*_2_ calculations. They then quantified the similarity across classes by measuring *D_inter_ – D_intra_* normalized by the maximum expected value of this difference. The normalization procedure is highly dependent on the type of system under consideration, and it might not be possible to calculate the denominator in many cases. PRED is self-normalizing and system agnostic, providing a consistent estimate of class separability for any dataset.

So far, we have computed *D*_1_ and *D*_2_ as the Euclidean distances between within-class and across-class values. In principle, one can use any distance measure in place of Euclidean distances for calculating PRED. For example, one can use Mahalanobis distance to account for different variabilities of the various dimensions of a response or Hamming distance to compare datasets with binary or categorical values. For temporal data, instead of binning the responses, one could use methods like the Victor-Purpura (Victor and Purpura, 1997, 1996) or the van Rossum (Rossum, 2001) distances to calculate the distance between spike trains. This flexibility in the choice of the distance metric may help in the future in optimizing PRED for different use cases.

## Materials and Methods

### Class-vector PRED

We generalized the definition of PRED from our previous work (Mittal et al., 2020) to all class-vector datasets. We considered all possible combinations of pairs of vectors and pairs of classes to calculate the PRED value. For each 2×2 matrix thus obtained, we computed two distances (**Figure 1a**): *D*_1_ = (*A*1 – *B*1)^2^ + (*A*2 – *B*2)^2^ is the sum of the squared Euclidean distances between the values to the same classes in different vectors; *D*_2_ = (*A*1 – *B*2)^2^ + (*A*2 – *B*1)^2^ is the sum of the squared distances between the values belonging to different classes in different vectors. We used the ratio 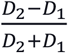 to estimate the PRED value in each 2 × 2 matrix. To obtain the final PRED value for a particular dataset, we first averaged the values over all class pairs before averaging over all vector pairs. Cases with missing data were ignored for the calculation of the mean. Note that in the calculations described here, the Euclidean distances can be easily calculated even if the values (*A*1, *B*1, *A*2, *B*2) are not numbers but are equal-sized vectors (see **Figure 3a** for an example). PRED ranges between 1 and −1, where 1 indicates that the vectors have identical values and patterns across classes, 0 indicates that the vectors have no similarity and have random patterns across the classes, and −1 indicates that the vectors have exactly opposite patterns across the classes.

### Class-sample PRED

We used a slightly modified method of calculating PRED (labeled ‘exhaustive PRED’) for class-sample datasets (**Figure 4a**). The calculation of *D*_1_ and *D*_2_ and the ratio 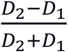 remained unchanged. The difference here lay in the creation of 2 × 2 matrices: for each pair of classes, any two samples (say, 1A and 1B) in class *i* could be combined with any two samples (say, 2A and 2B) in class *j*, to create two possible matrices, 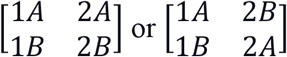. This results in a total of 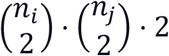 matrices for classes *i* and *j*, where *n_i_* = number of samples in class *i* and *n_j_* = number of samples in class *j* (see **Supplementary Figure 5a** for an example). We averaged the PRED values over all these matrices for each pair of classes and then computed the final PRED value by averaging over all class pairs.

### Other metrics for vector similarity in class-vector data

PRED was compared to 5 other metrics of vector similarity: Pearson’s correlation (PC), Cosine similarity (COS), Manhattan distance (MAN), Euclidean distance (EUC), and Chebyshev’s distance (CHEB). If the dataset included more than two vectors, each of the metrics was calculated over all possible pairs of vectors and then averaged. PC was computed using the corr function in MATLAB; while analyzing experimental datasets, any rows with incomplete data were removed. COS was as 1-cosine distance using the cosine option of the pdist function in MATLAB. The distance-based metrics MAN, EUC, and CHEB were calculated using the pdist function with the options cityblock, euclidean, and chebychev, respectively. Since the range of the distance-based metrics (MAN, EUC, and CHEB) was between 0 and ∞, we transformed these metrics using the negative exponential function *f*(*x*) = *e^−x^* which mapped the range to be between 1 and 0 such that a value close to 1 indicated a small distance (high similarity) between the vectors.

### Other metrics for class separability in class-sample data

PRED was compared to 5 other metrics of class separability: Euclidean template matching (ETM), Silhouette index (SIL), Davies-Bouldin index (DBI), Dunn’s index (DUNN), and Calinski-Harabasz index (CH). ETM is based on a simple algorithm for calculating classification accuracy (Stopfer et al., 2003). Briefly, a template was created for each class by averaging the values within the class, excluding the test sample. Next, for each sample in the dataset, the Euclidean distances between the sample and all the templates were calculated. If the smallest distance belongs to the template of the actual class of the sample, the sample was correctly classified and scored as 1 (if templates of n classes, including the actual class of the sample, had the same smallest distance, the score was set to 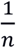). Otherwise, the score was set to 0. The average of the scores from all the samples was reported as the final value of ETM. ETM ranges between 0 and 1, where 1 denotes the highest level of class separability (every sample is correctly classified). We used a custom function written in MATLAB for calculating the ETM values. The Silhouette index compares the pairwise intra-class and interclass distances (Rousseeuw, 1987). It ranges between 1 and −1, where 1 indicates high separability. DBI is calculated as the ratio of within-class and between-class distances (Davies and Bouldin, 1979). It ranges from 0 to ∞, where 0 indicates high separability. CH measures the ratio of the average intra-class and inter-class variances (Caliński and Harabasz, 1974). It ranges between 0 and ∞, where a higher value indicates higher separability. SIL, DBI, and CH were calculated using the evalclusters function in MATLAB, with the options Silhouette, DaviesBouldin, and CalinskiHarabasz, respectively. DUNN calculates the ratio of the minimum inter-cluster distance to the maximum intra-cluster distance (Dunn, 1974). It ranges between 0 and ∞, where a higher value indicates high separability. We calculated the DUNN value using the indexDN function written by Julian Ramos for MATLAB.

### Simulations with clusters of points

To simulate a class-sample dataset, we first selected the class means uniformly distributed within an n-dimensional space [−1, 1]^*n*^. The samples were then drawn from a uniform distribution around the class mean such that the Euclidean distance between the sample and the class mean was ≤ *r*, where *r* denotes the cluster radius. Next, a random noise n-dimensional vector, drawn from 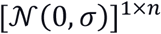, was added to each sample (see **Supplementary Figure 3f—h** for examples). Note that after the addition of noise, the samples no longer lay within [−1, 1]^*n*^ but, instead, within [−∞, ∞]^*n*^.

### Chance level

The chance level for each metric was calculated using datasets with no inherent similarity or separability. For the class-vector metrics, we simulated a dataset of 10 vectors and either 2 or 5 classes. Each value within the dataset was randomly drawn from a uniform distribution between −1 and 1, ensuring no structure within the classes or the vectors. The whole simulation was repeated 1000 times, and the vector similarity metrics were reported. For the class-sample metrics, we simulated a 2-dimensional clustered dataset with 10 samples and either 2 or 5 classes. The cluster radius was set to 0.05 for all the classes, and a big noise term randomly drawn from 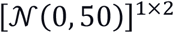 was added to simulate inseparable clusters. The whole simulation was repeated 1000 times, and the class separability metrics were reported.

### Dynamic range and variability

The dynamic range was defined as the range of noise levels in which a metric remains informative (i.e., does not saturate near the maximum or the minimum level). We simulated a dataset with increasing levels of noise (on a log scale). We measured the average value reported by the metric at the 5 lowest noise levels (as *μ*(*ν_l_*)) and at the 5 highest noise levels (as *μ*(*ν_h_*)) simulated. The absolute difference between these two values, |*μ*(*ν_l_*) – *μ*(*ν_h_*)|, was called the vertical range of the metric. For a metric whose value decreased with increasing noise, the left boundary of the dynamic range was taken as the lowest noise level at which the average value of the metric was lower than the value at the lowest noise level by at least 1% of the vertical range, i.e., *DR_l_* = min(*x*): *μ*(*x*) < *μ*(*ν_l_*) – 0.01 × |*μ*(*ν_l_*) – *μ*(*ν_h_*)|. The right boundary of the dynamic range was taken as the highest noise level at which the average metric value was greater than the value at the highest noise level tested plus 1% of the vertical range, i.e., *DR_h_* = max(*x*): *μ*(*x*) > *μ*(*ν_h_*) + 0.01 × |*μ*(*ν_l_*) – *μ*(*ν_h_*)|. The dynamic range was calculated as |*DR_h_* – *DR_l_*|.

The variability of the metric was defined as the standard deviation of the metric at the midpoint of the dynamic range divided by its vertical range, i.e.,

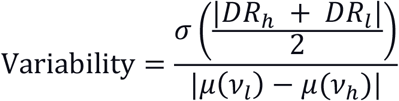

where *σ*(*x*) represents the standard deviation in the metric values at the noise level x. For the class-vector metrics, we simulated a dataset with 10 vectors and 2 classes. The mean response of each class was set to 2 and 4, respectively. The value for a class was randomly drawn from 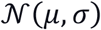, where *μ* is the class mean, *σ* = 10^*ν*^ and *ν* ∈ [−2, −1.9, −1.8,3] to simulate increasing noise levels on a log scale, covering 5 orders of magnitude. Each simulation was repeated 1000 times, and the resultant similarity was measured using each metric. We repeated the entire experiment with increasing base means, i.e., we added an integer value to the mean response of the classes. For example, adding 1 to the class means changed them from [2 4] to [3 5]. We simulated 11 such datasets by adding each of the integers in the range [0 10].

For class-sample datasets, we simulated a dataset with 2 classes, each with 10 samples. The response was set as a 2-dimensional vector. The class means were drawn from the 2-D space [−1 1]^2^ with a cluster radius of 0.05. The noise was drawn randomly from 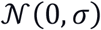, where *σ* = 10^*ν*^ and *ν* ∈ [−3, −2.9, −2.8,3] to simulate increasing noise levels (on a log scale) within the dataset. Each simulation was repeated 1000 times.

In the analysis where we added noise to the experimental data, we first calculated the mean response over all the different trials and odors (*m*). The noise (*ν*) was then added to each value of the data matrix as a percentage of this mean response with the values drawn from 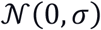, where *σ* = 10^*ν*^ X *m* X 0.01, *m* is the mean response, and *ν* ∈ [−1, −0.9, −0.8,…,4] is the noise level on a log scale.

### *Drosophila* olfactory behavior

We used a published dataset containing the behavioral preferences of 141 wild-type *Drosophila* for 3-octanol (OCT) versus 4-methylcyclohexanol (MCH) (Honegger et al., 2020). The behavior was quantified as a preference index obtained from a two-choice assay where the odors were presented, one on each port. A value above 0.5 indicated preference towards MCH while a value between 0 and 0.5 indicated preference towards OCT. The preferences were calculated for all the flies at two different time points, 24-hrs apart. We first calculated the individual-time (class-vector) PRED and Pearson’s correlation (PC) values over the entire dataset (**Figure 2a**). To compare the stability of the two metrics for incomplete data, we randomly sampled 70 out of 141 individuals from the dataset. We calculated the PRED and PC value for this subset, repeating the random sampling 20 times. We then calculated the coefficient of variation of each metric over these 20 random samplings. To check the validity of our results, we repeated this entire process 50 times and compared the coefficient of variation obtained from the two metrics.

### *Drosophila* population responses

To analyze the population level similarity in responses, we used a published dataset of calcium imaging responses of 37 glomeruli responding to 36 monomolecular odors (Badel et al., 2016). The glomeruli measured within the dataset were DM6, DM5, DM2, DM1, DM4, VM2, VM7d, VM7v, DA4L, DA2, DL1, DL5, D, DM3, DC2, VA6, DC3, DL4, DA3, DL3, DA1, VA1d, VA1v, VL2a, VL2p, VA5, VM4, VA7L, VA3, VA4, VA7m, VC2, VC1, VM3, VA2, VM1, and Dp1m. The odors used in the dataset were apple cider vinegar, mango mimic, broth, benzaldehyde, 2-methyl phenol, butanol, g-butyrolactone, methanoic acid, hexanoic acid, 1-octanol, acetophenone, vinegar mimic, 2,3-butanedione, pentanoic acid, 3-methylthio-1-propanol, 3-octanol, ethyl butyrate, 4-methylcyclohexanol, acetaldehyde, 2-pentanone, 2-oxopentanoic acid, hexyl acetate, isopentyl acetate, phenylethylamine, propionic acid, geosmin, ethyl acetate, *β*-citronellol, benzyl alcohol, linalool, 1-octen-3-ol, methyl salicylate, pentyl acetate, banana essence, 2-butanone, and 1-butanol. The dataset included the responses for 61 individuals (although not all individuals were measured for all odors) with around 4 trials each. For calculating the similarity within the individuals, we first averaged the responses over the trials. We then calculated the odor-individual (class-vector) PRED for each of the 37 different glomeruli separately (**Figure 3e**). Alternatively, we used the 37-length vectors as the values in the 61 (odor) X 36 (individual) matrix and calculated a single odor-individual (class-vector) PRED for these ‘population’ responses.

### Zebrafish olfactory behavior

We extracted the published data of seven behavioral responses of 10 wild-type Zebrafish in response to 18 different odors over 4 different trials from the raw data files provided by the authors (Kermen et al., 2020). The odors for which the response of the zebrafish was tested were food extract (food), histidine (his), nucleotides (nucl), methionine (met), phenylalanine (phe), cysteine (cys), arginine (arg), bile acids (bile), prostaglandin 2α (pgf2a), urea, ammonium (amo), putrescine (put), spermine (sperm), cadaverine (cad), chondroitin sulfate (cs), zebrafish blood (blood), zebrafish skin extract (skin), and artificial fish water (afw). The behaviors extracted were fish velocity, freezing behavior, vertical position in the arena, percentage of burst swimming, number of abrupt turns, number of horizontal swimming events, and number of vertical swimming events. We used custom scripts and MATLAB functions provided through personal correspondence by Dr. Florence Kermen to extract the data using the protocol described in the original paper (Kermen et al., 2020).

To characterize the individual-to-individual separability, we calculated the individual-time (class-sample) PRED value separately for each of the 18 odors. For each odor, the dataset included 10 classes (individuals) with 4 samples (trials) per class. The value of each sample was a 7-dimensional vector, representing the 7 behaviors (**Figure 7a**).

### *Drosophila* projection neuron responses with and without serotonin blockage

We obtained the published calcium imaging responses of 14 different projection neurons (PNs) from 18 different GCaMP6m wild-type flies and 7 a-methyl tryptophan (a-mw) fed flies to 12 different monomolecular odors (Honegger et al., 2020). The PNs in this dataset innervated DA1, DL3, DL1, DL5, DM3, DM6, DA2, DA4l, D, DM5, DM2, DM1, DM4, and DL4 glomeruli. The odors within the dataset were 3-octanol, 1-hexanol, ethyl-lactate, citronella, 2-heptanone, 1-pentanol, ethanol, geranyl-acetate, hexyl-acetate, 4-methylcyclohexanol, pentyl-acetate, 1-butanol. Each response was measured over 2 trials. We calculated individual-trial (class-sample) PRED separately for each PN-odor combination (**Figure 7c**).

### Locust and *Drosophila* electrophysiological recordings

We used published recordings of the response of bLN1 mushroom body output neurons in 6 different locusts responding to 6 different odors (Gupta and Stopfer, 2014). These electrophysiological responses were measured in awake locusts exposed to cyclohexanone, octanol, and hexanol in concentrations of 0.1% and 10% each. Each response consisted of 6-10 trials.

We also used the published responses of *Drosophila* PNs innervating 4 different glomeruli (VC4, DL2v, VM5v, VC3) to a set of 5 odors – benzaldehyde, 2-octanone, pentyl acetate, ethyl acetate, and ethyl butyrate (Shimizu and Stopfer, 2017) – although not all PNs were measured for all the odors. The response of each PN was measured in 2-6 individuals with approximately 6-10 trials per response.

For analyzing the odor-individual (class-vector) PRED with temporal responses, we extracted both the firing rate and the temporal response of the neurons for a period of 2-s after odor onset. The firing rate was calculated as the total number of spikes within the 2 second period from 2 to 4 seconds in the response minus the number of spikes in the 2 second period before odor onset, from 0 to 2 seconds in the response. The temporal response was similarly calculated in the 2 second period after odor onset divided into 10-bins of 200 ms each minus one-tenth the total number of spikes in the background response from 0 to 2 seconds. For calculating PRED and PC, we first averaged the responses over all the trials for each cell in the dataset (1 cell in the locust dataset and 4 cells in the *Drosophila* dataset). We then calculated the odor-individual (class-vector) PRED using both the firing rate (magnitude) and the temporal responses. PRED values were averaged over all pairs of odors for every pair of individuals.

In the experiments where we added noisy bins to the experimental datasets, we used the initial 10-bin vector of responses as the base dataset. For adding one noisy bin to the base dataset, we used the number of spikes obtained from 4 to 4.2 seconds minus the background response as the eleventh bin. Similarly, any extra noise bin extended the response period by 200 ms to a maximum of 4 seconds when 10 extra noise bins were added.

In the experiment, where we investigated the applicability of PRED to class-sample datasets, we used both the locust and the fly databases to calculate odor-trial (class-sample) PRED and compared it to the odor separability obtained from the other metrics. For each individual and cell in the dataset, we used the 2-bin (each bin of length 1 second) response vector to calculate the separability.

### Temporal response simulations

To simulate the temporal responses, we created a dataset with 2 classes and 10 vectors, where each response was a 10-bin vector. The base mean of each response bin within a class was randomly drawn from a uniform distribution in the range [1 3]. A random noise drawn from 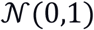 was added to each bin. A particular number of extra bins were appended to the vectors, with each new bin containing a value with a base mean of 0 and a noise drawn from 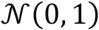. We compared the PRED and PC values with the number of extra bins ranging from 0 to 10. The entire simulation was repeated 100 times. To further emphasize the difference between the behaviors of PRED and PC, we repeated this entire simulation by generating extra bins that were exactly 0 (without any noise).

### Simulations with increasing numbers of classes or samples

We generated 2-dimensional clustered data with cluster means drawn from [−1 1]^1×2^ and cluster radius of 0.05. A small amount of noise drawn from 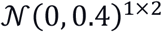 was added to each response in the dataset. For the simulations with increasing numbers of classes, we simulated two different datasets – one with 2 samples and the other with 10 samples. The number of classes ranged from 2 to 10. For the simulations with increasing numbers of samples, we used 2 classes. The number of samples was taken from [2, 4,…,20]. Each simulation was repeated 100 times.

For comparing ‘fast PRED’ with ‘exhaustive PRED’, we used the same dataset of 2 classes as described above but varied the number of samples from [2, 3,…, 25]. Each simulation was repeated 1000 times. The average value of PRED over all simulations was ~0.5. For each simulation and number of samples, we calculated the absolute difference between ‘fast PRED’ and ‘exhaustive PRED’ values. Finally, we reported the average difference over simulations divided by the average ‘exhaustive PRED’ value for the specified number of samples.

### *Drosophila* connectome data

We obtained the connectivity vectors of identified local horn neurons (LHNs) from Schlegel et al. (Schlegel et al., 2021) for 87 identified neurons in the FAFB and the Hemibrain databases. The dataset we used included 47 neurons from FAFB and 85 neurons from Hemibrain along with their connectivity to 57 unique antennal lobe glomeruli (D, DA1, DA2, DA3, DA4l, DA4m, DC1, DC2, DC3, DC4, DL1, DL2d, DL2v, DL3, DL4, DL5, DM1, DM2, DM3, DM4, DM5, DM6, DP1l, DP1m, V, VA1d, VA1v, VA2, VA3, VA4, VA5, VA6, VA7l, VA7m, VC1, VC2, VC3, VC4, VC5, VL1, VL2a, VL2p, VM1, VM2, VM3, VM4, VM5d, VM5v, VM6, VM7d, VM7v, VP1d, VP1l, VP1m, VP2, VP3, VP5). The LHNs were grouped into 49 ‘connectivity types,’ which were further grouped into 36 ‘cell types’, then 13 ‘tracts’, and finally 4 ‘regions’, based on their morphologies within the lateral horn (Frechter et al., 2019; Schlegel et al., 2021).

The full dataset consisted of unique connectivity types as classes and the two databases as vectors. The connectivity vector of each neuron within a connectivity type was averaged. Each cell within this matrix was a 57-length vector of averaged and normalized connectivity weights of the corresponding LHN to each glomerulus. We first calculated the connectivity type-database (class-vector) PRED value over this matrix to characterize the similarity of connections across databases. Next, we grouped this matrix based on each of the different hierarchy levels. We averaged the connectivity vectors over all connectivity types belonging to a group within a particular hierarchy to get a matrix with groups as columns and the databases as rows (**Figure 8a**). We then calculated the group-database (class-vector) PRED values for each hierarchy level based on cell type, tract, or region.

In the experiment where we characterized the separability of neurons across groups based on their connectivity to antennal lobe glomeruli, we constructed 4 different matrices with individual neurons (not averaged over connectivity types) as samples and the relevant group types as classes for the two databases separately (**Figure 8b**). We then calculated the group-neuron (class-sample) PRED for each matrix to characterize the separability of neural connectivity vectors across groups for each hierarchy level.

### Statistics

To compare a set of PRED values with the baseline (0) or a specific mean, we used a one-sample double-sided t-test. To compare the chance level of the metrics across classes, we used two-sample double-sided unpaired t-tests. For comparing the coefficient of variation obtained for PRED with those for PC, we used a two-sample double-sided paired t-test.

### Code availability

All the simulations and analyses were done using custom scripts coded in MATLAB (version r2020a). A modified version of the *gramm* plotting package (Morel, 2018) was used for all the figure plots. The source code for the simulations and analysis can be found at https://github.com/neuralsystems/PRED_analysis. The standalone versions of PRED function written in Python and MATLAB can be found at https://github.com/neuralsystems/PRED (the MATLAB version is also available on the MATLAB File Exchange).

## Competing interests

The authors declare no competing interests.

## Acknowledgments

We thank Kazumichi Shimizu and Mark Stopfer for sharing fly electrophysiology data and Hokto Kazama for sharing fly calcium imaging data. We thank Florence Kermen and Alexander Bates for helping us extract the zebrafish behavioral data and the connectomics data, respectively. We thank Arjit Kant Gupta for providing an initial version of the Python implementation for PRED. This work was supported by the DBT/Wellcome Trust India Alliance Fellowship [grant number IA/I/15/2/502091] awarded to N.G.; Cognitive Science Research Initiative of the Department of Science & Technology [DST/CSRI/2018/102] to N.G.; SERB Core Research Grant [CRG/2020/004719] to N.G.; a BBSRC grant [BB/S016031/1] to A.L.; and a Starting Grant from the European Research Council [639489] to A.L.

## Author Contributions (using CRediT format)

A.M.M. Conceptualization, Data Curation, Formal Analysis, Investigation, Methodology, Software, Visualization, Writing – original draft, Writing – review & editing

A.C.L. Conceptualization, Funding acquisition, Methodology, Writing–review and editing

N.G. Conceptualization, Formal analysis, Funding acquisition, Investigation, Project administration, Supervision, Visualization, Methodology, Writing–original draft, Writingreview and editing

## Notes

https://github.com/neuralsystems/PRED

### Competing Interest Statement

The authors have declared no competing interest.

