## Supplementary Figures for "Pairwise Relative Distance (PRED) is an intuitive and robust metric for assessing vector similarity and class separability"

### Supplementary Figure 1

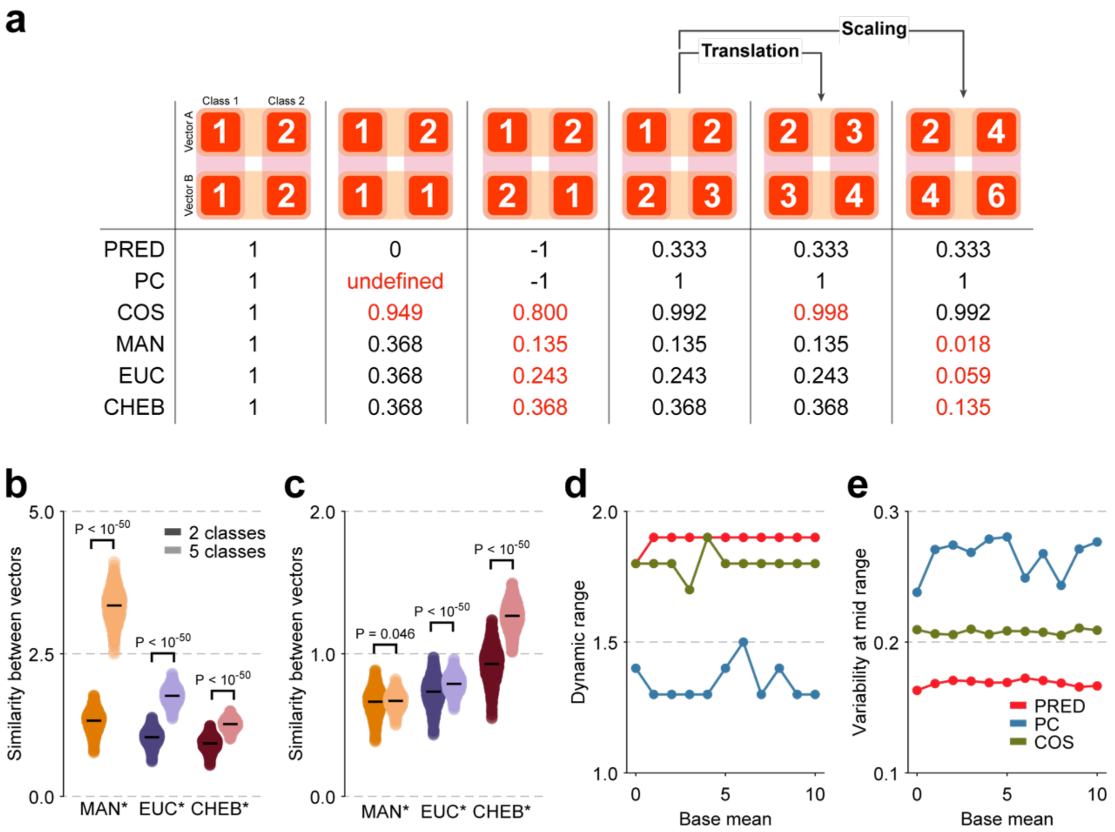


***Supplementary Figure 1: PRED is more stable to dataset modifications than other class-vector metrics***

**a** Examples comparing the values obtained from different metrics. Values in red represent less desirable outcomes, in our view, compared with PRED. **(i)—(vi)** In order from left to right: **(i)** Example with identical vectors for the 2 classes; **(ii)** Example where vector B has equal values for both the classes; **(iii)** Example where the two vectors have the opposite patterns across classes; **(iv)** A representative example with some similarity between the two vectors; **(v)** Example of a global translational modification where all the values in example iv are increased by 1; **(vi)** Example of a global scaling modification where all the values in example iv are multiplied by 2. **b** Change in the chance levels of the untransformed distance metrics with the number of classes in the dataset. Note that these untransformed metrics denote distance and not similarity. Hence, a high value is expected with completely random data. **c** Same as (**b**), but with MAN, EUC and CHEB values normalized by $n$, $n^{\frac{1}{2}}$ and $n^{\frac{1}{\infty}}$, respectively where $n$ denotes the number of classes (CHEB values are not affected as the normalization factor is 1). In (**b**) and (**c**), each point represents the value of the metric for a different random seed (n = 1000 simulations). Black horizontal line represents the mean. Error bars represent s.e.m. **d** Dynamic range remains consistent with increasing base mean of the classes for PRED, PC, and COS. **e** Variability remains consistent with the increasing base mean of the classes for PRED, PC, and COS.

### Supplementary Figure 2

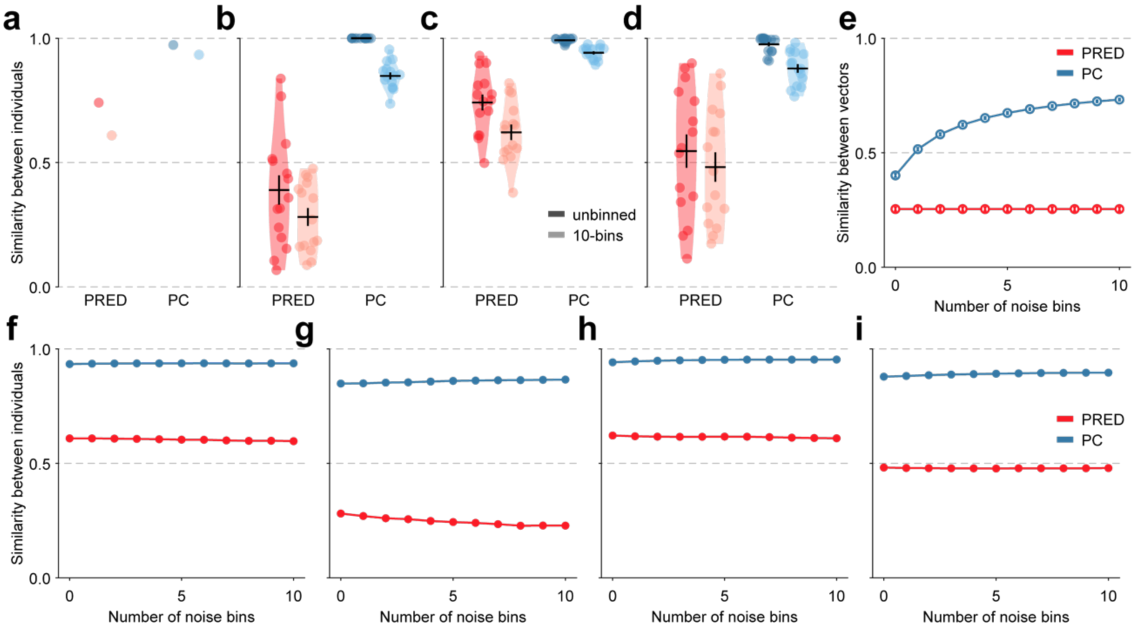


***Supplementary Figure 2: PRED remains informative even with a different dataset***

**a—d Across-**individual similarity when the neural response is quantified as a single number (darker colors) or as a 10-bin temporal vector (lighter colors) for four different neurons in the dataset. The data is taken from *Drosophila* projection neuron responses (Shimizu and Stopfer, 2017). Each point within the violin represents the similarity for a pair of individuals (a: n = 1 for both PRED and PC in both types of analyses, b: n = 15, c: n = 15, d: n = 15). Black horizontal lines represent the mean. Error bars represent s.e.m. in all panels. **e** Across-individual similarity as a function of the number of extra bins (containing noise) added to a 10-bin vector for simulated data with 2 odors and 10 individuals. The value in each extra bin was exactly 0. Open circles denote the mean over 100 different random simulations. Note that the similarity value reported by PC increases with the increasing number of bins, but PRED remains unaffected. **f—i** Across-individual similarity as a function of the number of extra bins (containing mostly noise) added to the original 10-bin vector for the same dataset as in **(a-d)**. Note that the similarity value reported by PC increases with the increasing number of bins.

### Supplementary Figure 3

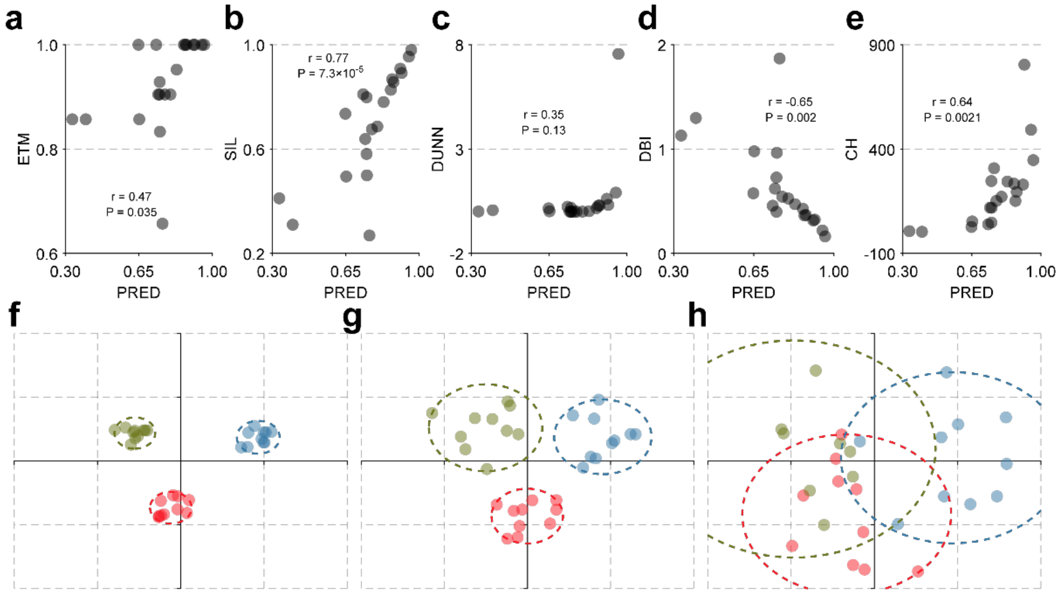


***Supplementary Figure 3: PRED can analyze datasets with clustered points***

**a—e** Odor separability measured using PRED compared to that measured using other commonly used metrics. Each point corresponds to one PN-individual combination in the dataset taken from *Drosophila* projection neuron responses (Shimizu and Stopfer, 2017) (n = 20). PRED: Pairwise relative distance, ETM: Euclidean template matching, SIL: Silhouette index, DUNN: Dunn's index, DBI: Davies-Bouldin index, CH: Calinski-Harabasz index. **f—h** Illustrative simulations with 3 classes (different colors) and 10 samples (points with the same color) showing the effect of increasing noise starting from low **(f)**, medium **(g)** to high **(h)** levels of noise on the class clusters. Colored circles represent the area covered by each class. Note that overlap between classes increases with increasing noise.

### Supplementary Figure 4

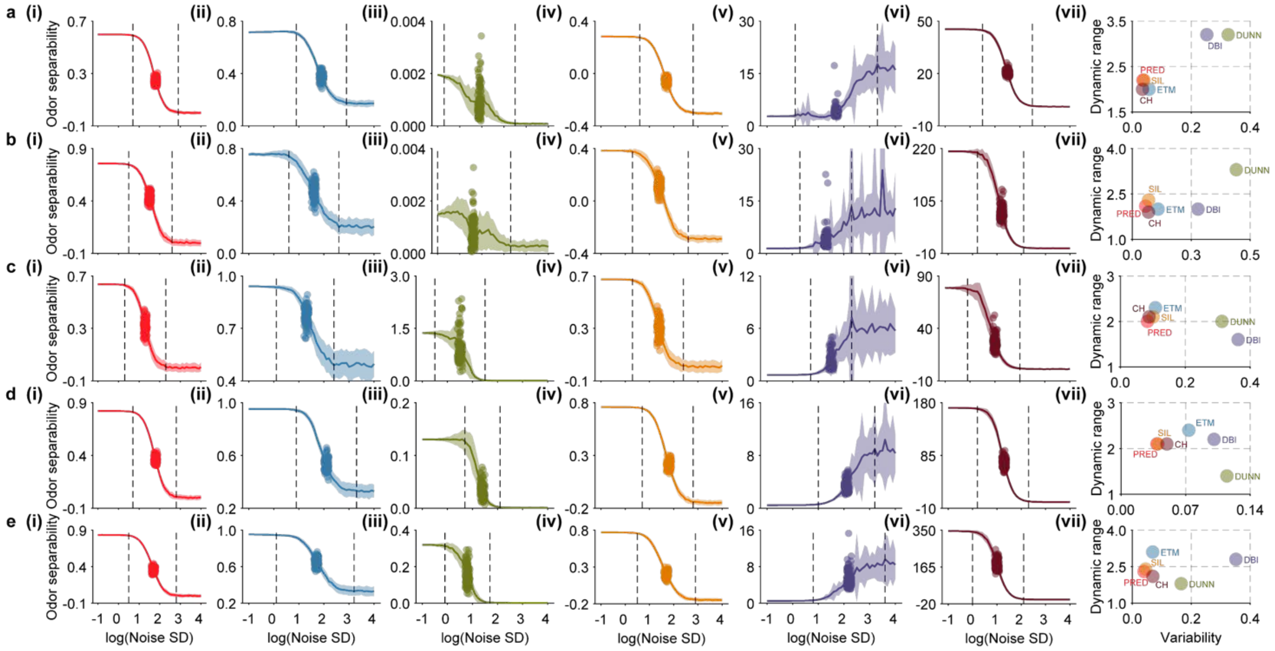


***Supplementary Figure 4: Comparison of metrics based on their robustness to noisy experimental data***

**a—e** Odor-separability reported by PRED (**i**), ETM (**ii**), DUNN (**iii**), SIL (**iv**), DBI (**v**), and CH (**vi**) with increasing levels of noise (shown on a log scale) added to experimental data obtained from locust bLN1 neurons (**a**) and from 4 different *Drosophila* PNs (**b-e**) (Gupta and Stopfer, 2014; Shimizu and Stopfer, 2017). **(i)—(vi)** The solid trace shows the mean values over all simulations for different noise levels (n = 1000 simulations). The shaded area represents 1 s.d. around the mean. The dashed vertical lines represent the boundaries of the dynamic range. Each point represents a different random simulation at the noise level corresponding to the mid-point of the dynamic range. **(vii)** The dynamic range and the variability at the mid-point of the dynamic range are shown for each metric. PRED consistently showed a large dynamic range and low variability.

### Supplementary Figure 5

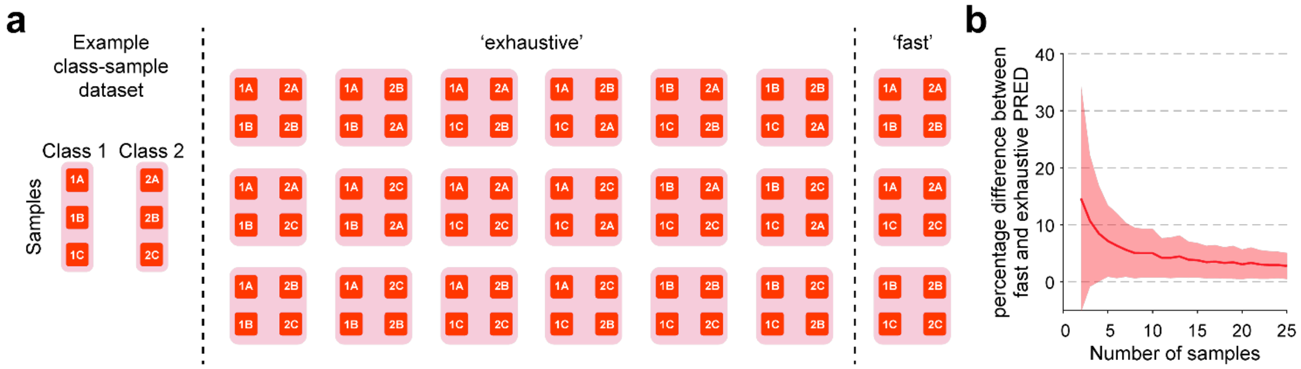


***Supplementary Figure 5: Error rate of 'fast PRED' reduces with the number of samples in class-sample datasets***

**a** Example illustration of all the pairwise combinations of classes and samples on which PRED computation is performed while calculating class-sample PRED with the 'exhaustive' or the 'fast' method. For this dataset with 2 classes, each having 3 samples, the number of computations required for 'fast' PRED is $\binom{3}{2} \cdot \binom{2}{2} = 3$ whereas for 'exhaustive' PRED it is $\binom{3}{2} \cdot \binom{3}{2} \cdot 2 = 18$. The final PRED value is obtained by averaging over all computations within each method. **b** For a simulated dataset of 2 classes and the indicated number of samples per class, the absolute difference between the values obtained from 'fast PRED' and 'exhaustive PRED' is shown as a percentage of the average 'exhaustive PRED' value over all iterations. Note that 'fast PRED' exhibits an error of only ~3% if the dataset contains more than 15 samples per class. The dark red line represents the mean of 1000 different iterations at each point. The shading represents s.d. around the mean.
